# Recombinase Polymerase Amplification of Forensic Short Tandem Repeat Loci

**DOI:** 10.1101/2025.07.01.662531

**Authors:** Sonja Škevin, Liesl De Keyzer, Lynn De Waele, Olivier Tytgat, David Van Hoofstat, Dieter Deforce, Filip Van Nieuwerburgh

**Affiliations:** Laboratory of Pharmaceutical Biotechnology, Ghent University, Ottergemsesteenweg 460, Gent, 9000, Belgium; Eurofins Forensics Belgium NV, Lieven Bauwensstraat 6, 8200, Brugge, Belgium

**Keywords:** Recombinase polymerase amplification, short tandem repeats, microsatellites, forensic genetics, capillary electrophoresis, massively parallel sequencing, nanopore sequencing

## Abstract

Short tandem repeats (STRs) are highly polymorphic repetitive DNA sequences extensively used in forensic science for identification of individuals. STR genotyping is usually performed by capillary electrophoresis (CE) or next-generation sequencing (NGS) in centralized laboratories. However, there is an increasing need for a low cost, portable and rapid STR genotyping method. Multiple methods for miniaturization have been explored, all relying on polymerase chain reaction (PCR) for generating amplicons. PCR requires precise thermal cycling, which complicates the design of the STR genotyping microfluidic device. Recombinase Polymerase Amplification (RPA) is an isothermal DNA amplification method that operates between 37°C and 42°C and completes within 40 minutes. This, along with the robustness of reagents and reduced stutter rate compared to PCR makes RPA a suitable candidate for implementation in an STR genotyping microfluidics device as well as a part of the established STR genotyping work flows. In this proof-of-concept study, we evaluate RPA assay for amplification of forensically relevant STR loci. Thirteen core STR loci of the Combined DNA Index System (CODIS) were amplified using RPA in both singleplex and multiplex formats. The amplicons were then analyzed using three different methods: CE, Illumina and Oxford Nanopore Technologies (ONT). A subset of 5 loci was used for CE analysis. CE, Illumina and ONT sequencing of singleplex RPA each resulted in complete and correct STR profiles across all samples. Sensitivity assessment demonstrated that complete and correct genotypes were achieved with DNA inputs of 62 pg and above for all but locus D8S1179. Attempts at multiplex RPA amplification resulted in incomplete or incorrect STR profiles. This outcome highlights a challenge in adapting RPA for simultaneous amplification of multiple STR loci, which is a standard requirement in forensic DNA profiling.

## Introduction

Short tandem repeats (STRs), also known as microsatellites, consist of deoxyribonucleotide acid (DNA) sequences ranging from 1 to 6 base pairs, repeated multiple times in a head-to-tail manner. These sequences comprise approximately 3% of the human genome^1^ and exhibit mutation rates 10000 to 100000 times higher than the average across the genome. While initially considered neutral genetic markers, STR expansions have since been linked to numerous neurological and developmental disorders. Beyond their pathological implications, STRs play a role in DNA replication and repair, chromatin organization, and gene expression regulation^2^. Given their high polymorphism rate, STRs are extensively employed as genetic markers in humans, animals, and plants for various applications, including conservation, population genetics, kinship analysis, forensic DNA fingerprinting, and cancer diagnostics^3^.

The gold standard for STR genotyping involves amplicon sizing using capillary electrophoresis (CE). CE analysis is, since recently, complemented by the introduction of massively parallel sequencing (MPS), which offers distinct benefits, being an increased throughput, isoallele typing and single nucleotide polymorphism (SNP) detection^4^.

Even though both CE and MPS are established and reliable STR genotyping methods, there is an increasing demand for portable, rapid and affordable forensic genotyping^5^. Two main approaches to achieving speed and portability are currently pursued: a commercially available, portable CE device and a microfluidic-based STR genotyping approach^6^. Portable devices currently on the market are mostly CE-based, such as ANDE Rapid DNA (ANDE Corporations, Longmont, CO, USA) and RapidHIT™ ID System (Applied Biosystems, Thermo Fisher, Waltham, MA, USA). Although rapid and reliable, both devices come with an upfront cost of more than 200 000 USD. The price of the device, along with the high reagent cost, can be too high for many small, resource-scarce communities. This results in a significant backlog leading to delays in justice and closure for the victims^7^.

In addition to the established STR genotyping methods and aforementioned portable solutions, there is a growing body of research on STR genotyping using long read sequencing. Specifically, multiple recent studies have demonstrated the improvement of Oxford Nanopore Technologies (ONT) sequencing accuracy in STR genotyping^8,9,10^. One of ONTs products is a cost-effective, portable, universal serial bus (USB) powered MinION device, which has been repeatedly tested for on-site applications^11,12^. Therefore, MinION is another alternative for portable STR genotyping that does not come with a high upfront cost.

All of the described methods rely on PCR, which requires precise thermal cycling, which poses an additional obstacle for miniaturization. There are several isothermal amplification methods that are being researched with the purpose of circumventing the thermal cycling requirements of PCR, one of which is recombinase polymerase amplification (RPA). RPA is an isothermal DNA amplification method invented in 2006^13^ and it has primarily been researched in infectious disease diagnostics as a real-time pathogen detection method. RPA operates optimally in a temperature range between 37 and 42°C, but can function at room temperature, albeit with a reduced efficacy^14^. The RPA reaction is completed in up to 40 minutes and has a sensitivity on-par to PCR^15^. An additional benefit of the RPA assay is the robustness of the reagents, with no reduction in performance after storage at temperatures up to 45°C, for multiple weeks^16,17^. It has also been proven that RPA reduces stutter in cancer-related microsatellites compared to PCR^3^, a valuable characteristic in interpretation of low-input forensic samples and mixture samples.

The goal of this proof-of concept study is to explore the ability of RPA to amplify forensic STR loci with an envisioned application in either a portable microfluidic STR genotyping device, or as a part of an established forensic genotyping workflow. We used CE and MPS to explore the RPA amplicons of STR loci in order to address several key questions that arise when considering RPA for forensic STR amplification. These include the feasibility of working with fluorescently labeled markers, specificity, sensitivity, as well as speed and multiplexing capability. For this purpose, thirteen core STR loci of the Combined DNA Index System (CODIS) were amplified using RPA in singleplex and multiplexes. The amplicons of all thirteen loci were then analyzed using Illumina and Oxford Nanopore Technologies (ONT). A subset of 5 loci was used for CE analysis.

## Results

### RPA assay results

The success of the RPA assay was first verified on a 2% agarose gel (**Figure 1**). The singleplex RPA product displayed distinct bands at the anticipated amplicon lengths. Besides the amplicon bands, a band approximately 100 bp longer than the amplicon band is visible for several loci. Additionally, a smear is visible for most loci. A high level of primer artifacts is visible in the negative control lane, manifesting as a smear with equidistant bands. Thirteenplex RPA products exhibited a smeared profile with an unclear differentiation between bands (**Supplementary Fig. S1**).

**Figure 1.**
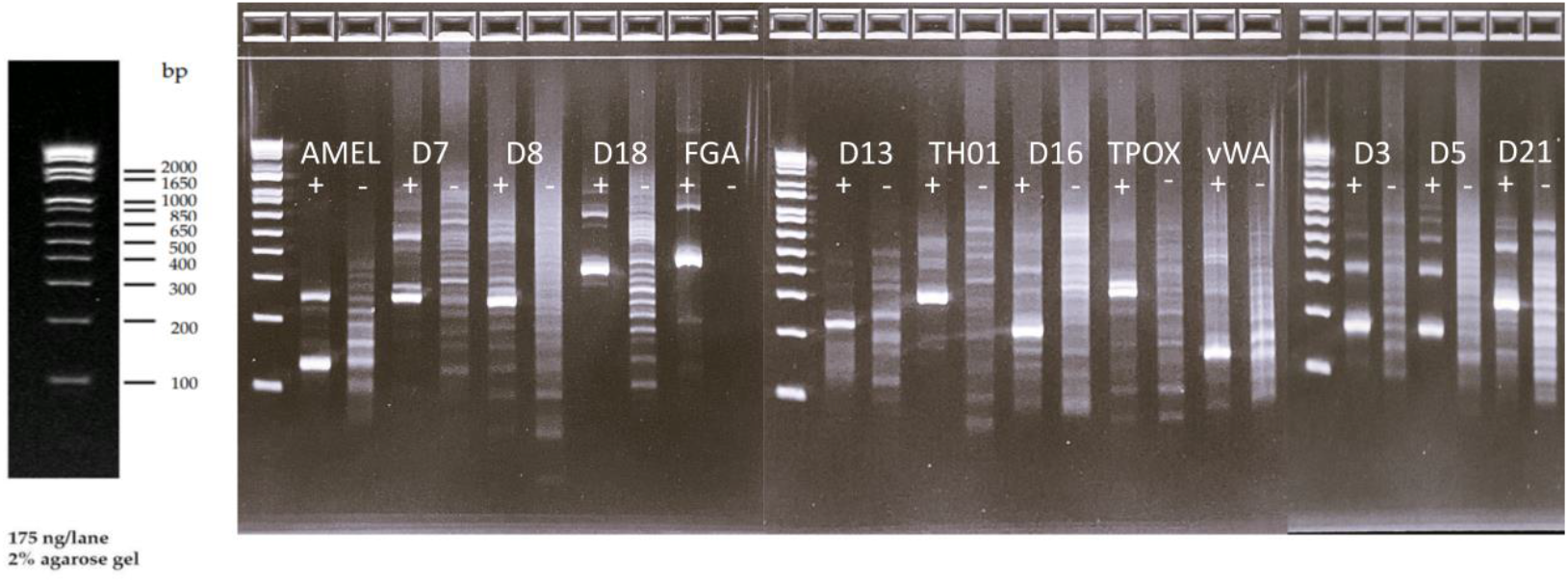
Singleplex RPA product of sample 3 (“+” labeled lanes) on a 2% agarose gel. Lanes labeled with “-” contain negative controls. A marker indicating band lengths is shown on the leftmost side of the figure.

### Capillary electrophoresis genotyping accuracy

All profiles obtained by CE were concordant with the profiles obtained by the benchmarking method for all eight samples tested. **Figure 2** presents an electropherogram of sample 4, while the electropherograms for the remaining samples amplified in singleplex can be found in the **Supplementary File**. Electropherograms show distinct peaks corresponding only to the expected amplicon length, without discernible noise or artifact peaks. The presence and intensity of stutter products, a common artifact in PCR-based STR analysis, varied among the different loci amplified by RPA. Notably, no detectable stutter products were observed for the STR loci TH01 and D8S1179 across all samples analyzed. Small stutter peaks were visible in the electropherograms for loci D18S51 and D21S11 in six tested samples. These stutter peaks were below the level of quantification set in the CE analysis software, and therefore have no impact on genotyping accuracy.

**Figure 2.**
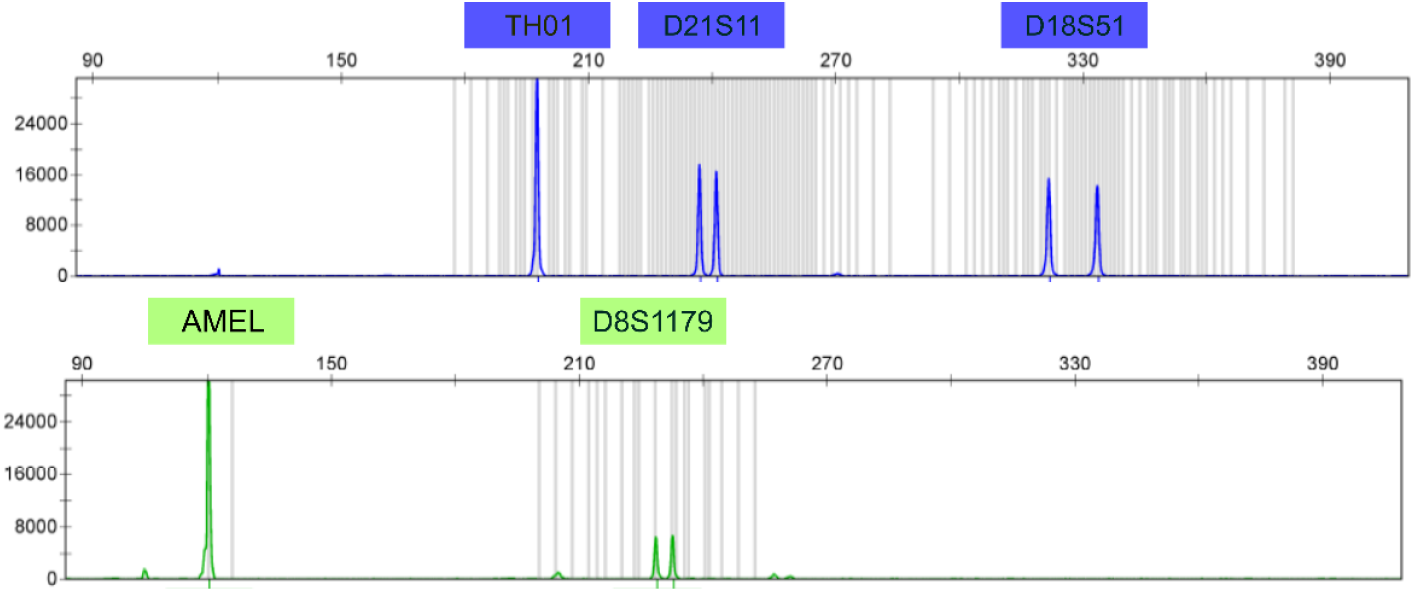
Capillary electrophoresis electropherogram of the pooled singleplex RPA products of sample 4, generated using a DNA input of 1 ng. Small, off-ladder peaks are visible within the size ranges of loci Amelogenin, D8S1179 and D21S11.

Allelic balance was assessed by calculating the ratio between the peak heights of both alleles (RFU peak 1/RFU peak 2) in the heterozygous samples across the loci. The average allelic balance was consistently between 0.75 and 1 in all samples and loci except for Amelogenin (**Table 1**).

**Table 1.** Average allelic balance (Avg. AB) and average peak height (Avg. PH), with corresponding standard deviations, for heterozygous loci amplified in RPA singleplex reactions and analyzed by capillary electrophoresis. Allelic balance is expressed as the ratio of the lower to the higher peak height of the two alleles in heterozygous samples. The column ‘n’ indicates the number of heterozygous samples observed per locus.

| Locus | Avg. AB | SD (AB) | Avg.<br>PH/RFU | SD<br>(PH)/RFU | Relative<br>SD (PH) | n |
| --- | --- | --- | --- | --- | --- | --- |
| Amel | 0.45 | 0.016 | 18272 | 9957 | 54.4% | 2 |
| D8S1179 | 0.72 | 0.26 | 7597 | 6601 | 86.9% | 7 |
| D18S51 | 0.76 | 0.22 | 10227 | 5561 | 54.4% | 8 |
| D21S11 | 0.82 | 0.20 | 16329 | 6042 | 37.0% | 6 |
| TH01 | 0.72 | 0.21 | 11567 | 8198 | 70.9% | 7 |

Analysis of fiveplex RPA products by CE resulted in incomplete or incorrect STR profiles across all samples. The performance varied considerably among the different loci: D21S11 showed the highest success rate, with correct allele peaks observed in 6 out of 8 samples. Amelogenin, D8S1179 and D18S51 each displayed correct allele peaks in 5 out of 8 samples. Locus TH01 shows faint, but correct peaks in only one sample. The electropherograms of fiveplex RPA products exhibited variability in peak intensity both between samples and among loci within samples. Additionally, numerous off-ladder peaks were observed, possibly attributable to non-specific amplification and primer-dimer artifacts. Electropherograms of RPA fiveplex samples, are available in the **Supplementary File**.

### Sensitivity assessment

Full STR profiles were consistently obtained with DNA inputs of 250 pg and 125 pg in both experimental replicates. One of the two 62 pg replicates showed allelic dropout at the D8S1179 locus, while the other replicate produced a complete profile. Input amounts of 500 pg and 31 pg resulted in D8S1179 peaks that were below the analytical threshold of the analysis software. **Fig. 3** shows the results of the sensitivity assessment.

**Figure 3.**
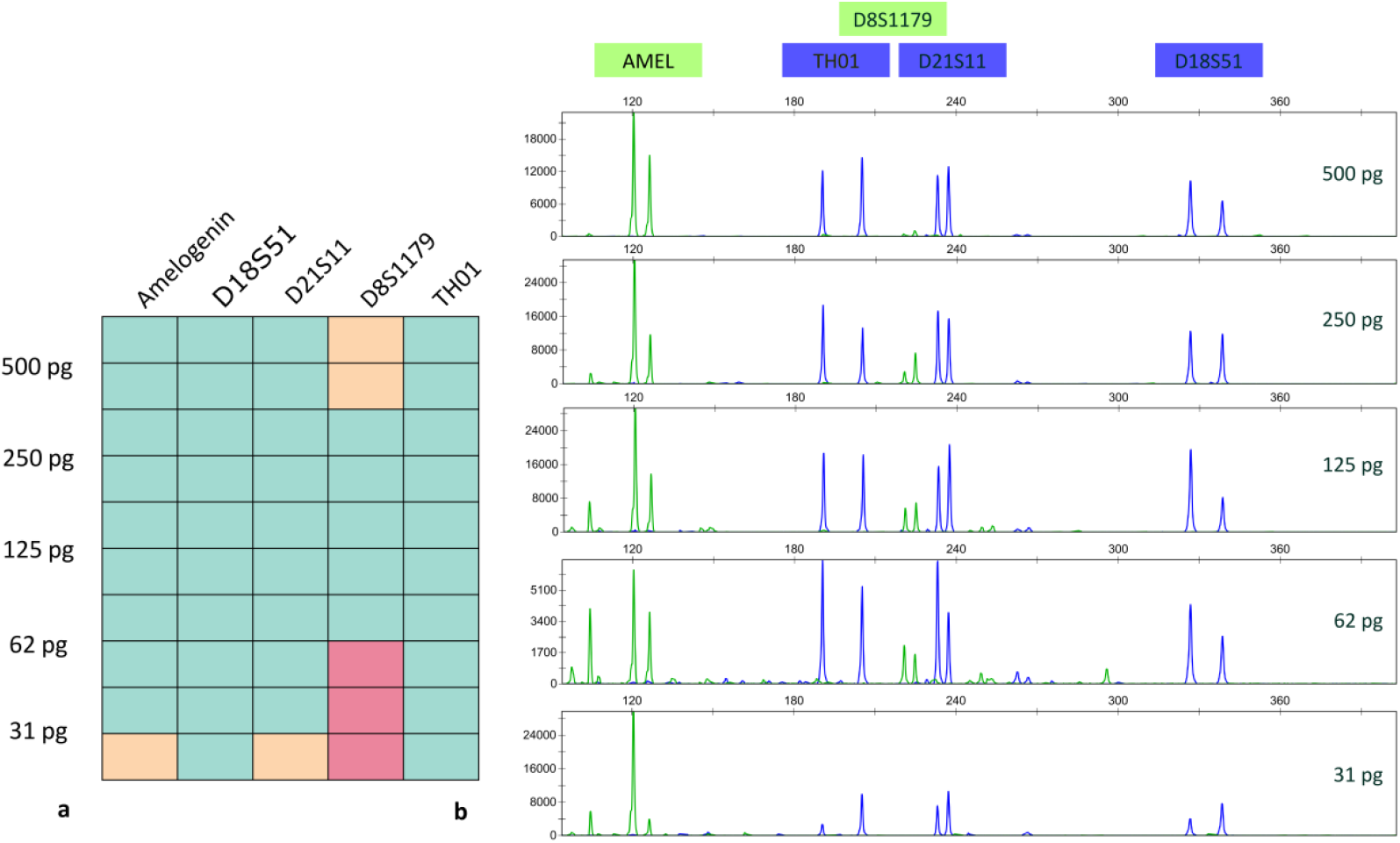
Sensitivity assessment of RPA singleplex using a range of DNA input amounts (31 – 500 pg) of sample 2, performed in duplicate. **a**. Genotyping concordance table: green fields indicate concordant genotypes for both alleles, orange fields indicate a single dropout, and red fields represent a double dropout. **b**. Electropherograms of the first replicate of singleplex RPA products using different DNA input amounts.

### Sequencing and genotyping results of RPA singleplex

On average, 164193 (SD=16080) reads were generated per singleplex sample during the Illumina sequencing run. Of these, 94.4% of generated pairs were successfully merged using FLASH tools. An average of 61.41% merged reads were assigned to a locus. Of the reads assigned to a locus, over 90% were uniquely mapped to an allele. The fraction of mapped reads mapping to a true allele was over 94% in all loci (**Table 2**).

**Table 2.** Number of reads mapped per locus and the percentages of those reads that map to a true allele in singleplex RPA setup after Illumina and ONT sequencing.

|  | Illumina |  | ONT |  |
| --- | --- | --- | --- | --- |
| | Reads mapping to true allele (%) | $\Sigma_{\text{reads per locus}}$ | Reads mapping to true allele (%) | $\Sigma_{\text{reads per locus}}$ |
| AMEL | 98.43 | 30377 | 90.48 | 435976 |
| D13S317 | 97.02 | 28523 | 90.77 | 1065401 |
| D16S539 | 98.75 | 52668 | 95.68 | 2682183 |
| D18S51 | 94.42 | 39748 | 76.50 | 3731515 |
| D21S11 | 95.13 | 30918 | 83.96 | 1117293 |
| D3S1358 | 95.83 | 28614 | 86.28 | 1149911 |
| D5S818 | 95.78 | 23193 | 76.16 | 950446 |
| D7S820 | 97.92 | 35907 | 79.07 | 1440724 |
| D8S1179 | 97.58 | 44429 | 90.14 | 11543 |
| FGA | 95.15 | 33437 | 60.43 | 3677815 |
| TH01 | 99.57 | 19358 | 96.37 | 1390858 |
| TPOX | 99.43 | 8568 | 95.11 | 712143 |
| vWA | 94.59 | 52853 | 86.79 | 922422 |

Concordant STR profiles were obtained for all samples and loci in the singleplex setup (**Figure 4**). The allelic balance (AB) in heterozygous samples was between 0.7 and 0.85 for all loci except FGA amplicons sequenced by ONT (**Table 3**).

**Table 3.** Average allelic balance (AB) of heterozygous singleplex samples. AB is represented as the number of reads mapping to the allele with a lower count, divided by the number of reads mapping to the higher count allele. The resulting ratio was averaged across the heterozygous loci of 4 samples amplified in singleplex. The number of heterozygous loci across the different samples is indicated in column n.

| Locus | Illumina |  | ONT |  | n |
| --- | --- | --- | --- | --- | --- |
|  | Avg. AB | SD | Avg. AB | SD |  |
| AMEL | 0.53 | NA | 0.68 | NA | 1 |
| D13S317 | 0.75 | NA | 0.77 | NA | 1 |
| D16S539 | 0.76 | 0.18 | 0.78 | 0.16 | 3 |
| D18S51 | 0.74 | 0.09 | 0.76 | 0.12 | 4 |
| D21S11 | 0.76 | 0.10 | 0.77 | 0.14 | 3 |
| D3S1358 | 0.73 | 0.09 | 0.80 | 0.04 | 3 |
| D5S818 | 0.74 | 0.11 | 0.70 | 0.06 | 3 |
| D7S820 | 0.70 | 0.16 | 0.76 | 0.14 | 3 |
| D8S1179 | 0.79 | 0.22 | 0.68 | 0.10 | 4 |
| FGA | 0.84 | 0.17 | 0.83 | 0.19 | 4 |
| TH01 | 0.83 | 0.06 | 0.88 | 0.09 | 3 |
| TPOX | 0.72 | 0.23 | 0.70 | 0.21 | 3 |
| vWA | 0.83 | 0.08 | 0.91 | 0.003 | 2 |

On average, 8705546 (SD=824405) passed-filter reads were generated per singleplex sample during the ONT sequencing run. Following rebasecalling, on average 63.2% of the passed-filter reads were assigned to a locus. More than 92% of the assigned reads were uniquely mapped to an allele. The fraction of reads mapping to a true allele ranged from 60.43% to 96.37% between loci. The percentages of reads mapping to a true allele are shown in **Table 2**. The STR profiles of samples amplified in singleplex were concordant with those obtained by benchmarking across all samples and loci. Similarly to the Illumina sequencing data, the allelic balance in heterozygous loci as measured with ONT sequencing, was between 0.7 and 0.85 for all loci, except for FGA (**Table 3**).

### Multiplex RPA sequencing and genotyping

Illumina sequencing of the sixplex and thirteenplex samples resulted in on average 313020 (SD=25594) and 277158 (SD=20404) reads, respectively, of which over 94% were successfully merged by FLASH tool. On average, 53.66% and 27.40% of the merged reads were assigned to a locus in the sixplex and thirteenplex setups, respectively. Of the assigned reads, 93% and 89% were uniquely mapped to an allele in sixplex and thirteenplex, respectively. In the sixplex setup, only Amelogenin was incorrectly genotyped as X:X in sample 2, while the true genotype was X:Y. In the thirteenplex setup, only loci Amelogenin, D13S317 and D5S818 had sufficient coverage to establish a genotype in all samples, leading to correct genotypes for D13S317 and D5S818 and imbalanced results in the male samples for Amelogenin (**Fig. 4**).

**Figure 4.**
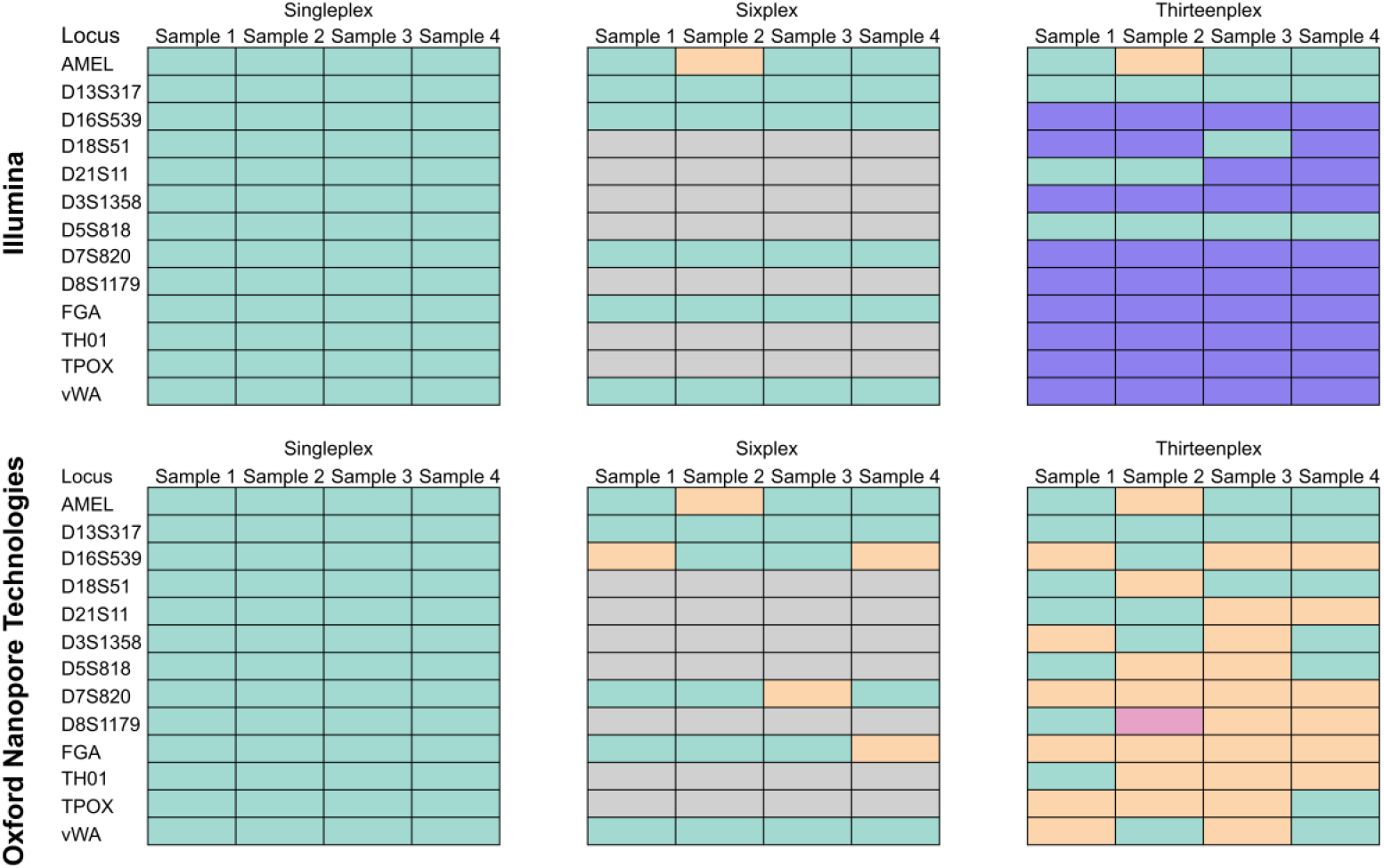
Genotyping results of Illumina and ONT sequencing of singleplex, sixplex and thirteenplex setups. Green fields indicate concordant genotyping for both alleles, orange fields indicate one non-concordant allele and pink field indicates both alleles were non-concordant. Purple fields indicate the coverage is too low to establish a reliable genotype and grey indicates the loci that were not included in the RPA sixplex.

To assess the amplification bias within the multiplex setups, the locus representation was calculated for each locus as a ratio of the number of reads assigned to the selected locus over the sum of all reads assigned to a locus. Graphic representation of the amplification bias for data generated via Illumina sequencing for all samples amplified in both sixplex and thirteenplex setups are shown in **Figure 5**. The most abundant locus in both setups was Amelogenin, which is also the shortest locus. In the sixplex setup, the second most abundant locus was vWA, but its representation drops from 32.5% to 0.27% when transitioning from sixplex to thirteenplex setup.

**Figure 5.**
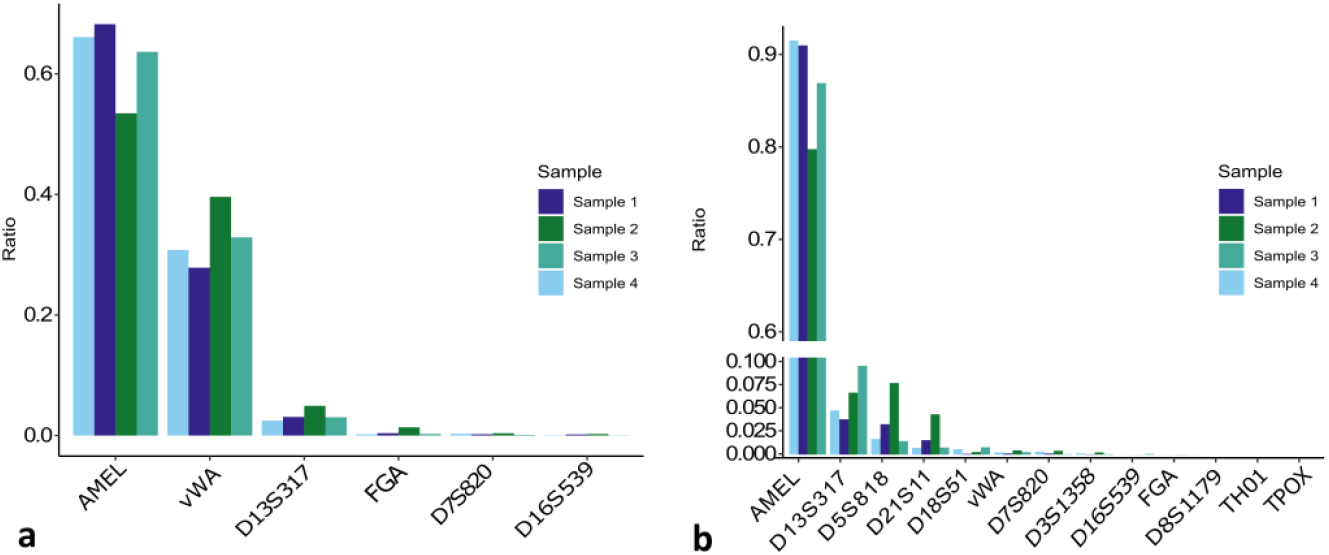
Amplification bias in RPA multiplexes, as measured by Illumina sequencing. The Y-axis shows the ratio of reads assigned to a certain locus divided by the sum of all reads assigned to any locus in **a**. Sixplex setup and **b**. Thirteenplex setup. Notably, the analysis performed with ONT sequencing produced very similar results (**Supplementary Fig. S2**).

ONT sequencing metrics, as well as the locus amplification bias diagrams of sixplex and thirteenplex can be found in the **Supplementary File**. Nanopore sequencing of the sixplex RPA product resulted in correct genotypes for loci D13S317 and vWA across all samples, while the loci Amelogenin, FGA and D7S820 were genotyped incorrectly in one of the samples and locus D16S539 was genotyped incorrectly in two samples. The thirteenplex RPA product exhibited higher error rates, with only locus D13S317 genotyped correctly across all samples (**Figure 4)**.

## Discussion

### STR genotyping of singleplex RPA amplicons

Distinct bands at anticipated lengths on an agarose gel of singleplex RPA product demonstrate that STR loci were successfully amplified using RPA. The presence of artifacts, including longer bands and smears on the agarose gels, could raise a concern regarding the specificity of the RPA product as non-specific products could complicate interpretation of results, especially in cases with limited or degraded DNA. However, concordant genotyping of all singleplex samples across all methods of analysis tested in this study prove that the artifacts seen on the gel have no detrimental effect on the genotyping results, as both the CE profiles and sequencing results show the correct genotypes. It is expected for the number of dropouts to increase when lower DNA input amounts were used, as it has been regularly reported for input amounts under 100 pg ^1,18^. Accordingly, we report dropouts in loci Amelogenin and D21S11 with an input of 31 pg and in D8S1179 for input amounts of 500 pg, 62 pg and 31 pg. Additionally, locus D8S1179 shows the lowest average peak height and highest peak height variability, which implies the lack of robustness of the D8S1179 assay (**Table 1**). However, only two replicates of one sample were included in this sensitivity study, therefore the dropouts may have occurred stochastically. More samples and replicates should be analyzed in order to have a robust assessment of the assay sensitivity. Extensive optimization for each locus specifically was not performed for this proof-of-concept study. Changes in primer design should be implemented in order to increase the efficiency and robustness of the D8S1179 assay. Nevertheless, this preliminary singleplex RPA sensitivity assessment gives an indication of RPA’s sensitivity when amplifying STR loci. The resulting sensitivity is comparable to that of the established STR genotyping kits, such as AmpFlSTR™ Identifiler™ Plus, which consistently yields full profiles at >125 pg ^19^. Average AB across both sequencing methods and CE was more than 0.7 (SD=0.2) in all loci amplified in singleplex except Amelogenin (0,45-0,78) (Table 1 & 3). AB in widely used commercial kits, as evaluated in the 2016 study^20^, is around 60%, which is comparable to the AB achieved using RPA singleplex. In profiles generated by Illumina sequencing, proportion of the reads mapping to the Y allele of Amelogenin locus was only 53% of the X allele read count. Similarly, CE profiles demonstrated a below average level of balance between X and Y peak height, with average Y allele peak height being 45% of X allele peak height. Further research is needed to establish the causes of the imbalance between the X and Y allele in Amelogenin locus when amplified using RPA. Nevertheless, in the electropherograms, the Y peak is still well above the analytical threshold and can therefore confidently be called as present.

### Multiplexing of the RPA assay

In the sixplex setup, Illumina sequencing yields correct genotypes for all loci except Amelogenin in sample 2, where incorrect genotyping stems from the allelic imbalance resulting in Y allele dropout. The allelic balance between Y and X alleles in female samples were 0.1% and 0.003% for sixplex and thirteenplex, respectively, compared to 9.6% and 19.3% in the male sample. This suggests that a different allelic balance threshold should be considered for calling a male sample for Amelogenin in multiplex RPA setups.

Coverage for samples amplified in sixplex was above the set coverage threshold for all loci, although a considerable amount of amplification bias was observed. In the sixplex setup, the Amelogenin and vWA loci accounted for an average of 96.7% of reads, while the least represented locus, D16S539, was represented by only 0.1% of the reads, which still satisfied the read count threshold. In the thirteenplex setup, a shift occurred. The fraction of vWA reads dropped from 32.5% to 0.2%, and the fraction of FGA reads dropped from 0.5%, which still provided sufficient coverage, to 0.009%. These changes in amplification levels are likely due to primer interactions introduced by the addition of primer sets to the singleplex and sixplex, as all loci amplified well in the singleplex and satisfactory in sixplex. Since different loci are used in the fiveplex panel than in the sixplex, the interactions between primers of different is likely the reason dropouts and incorrect peaks occur in the CE profiles of the fiveplex samples. The absence of denaturation and annealing steps in RPA, combined with increased total oligonucleotide concentration in the thirteenplex, promotes the amplification of primer dimers which competes with the amplification of the target sequence. These findings are consistent with the observed off-ladder peaks that were seen on the electropherograms of fiveplex samples (**Supplementary File**).

In this proof-of-concept study, extensive optimization and validation of the RPA multiplexes was not conducted. However, adequate coverage and high genotyping concordance of the sixplex reaction prove the potential of RPA for multiplex amplification of forensic STR loci. At the moment, and to the best of our knowledge, sixplex described in our study is the multiplex with the highest number of loci amplified at once using RPA. Most published RPA multiplexes involve no more than four primer sets at once^21^ and are primarily focused on pathogen detection in animals, humans, food and plants^15^. Therefore, literature reporting optimal RPA multiplex conditions is not yet available. Optimizations of RPA multiplex encompasses further adjustment of relative primer concentrations, as well as their length and sequence. Reaction temperature and duration, MgOAc and dNTP concentrations are also variables that have a potential in improving the RPA multiplex performance.

While our findings suggest that PCR could potentially be replaced by RPA in current centralized forensic laboratories, we acknowledge that there is no immediate demand for such a transition. Implementing RPA as a replacement would require multiple, extensive validation studies, which fall outside the scope of this proof-of-concept research. We believe in the potential for integrating RPA into a microfluidic STR genotyping device, but we are aware this remains a theoretical possibility. Developing a market-ready, lab-on-a-chip device for forensic RPA would require significant investment and effort, well beyond the scope of this proof-of-concept study. Nonetheless, we hope our manuscript inspires future initiatives to pursue this vision. Given the focus of our study, it would be inappropriate to compare our findings to state-of-the-art portable solutions such as ANDE or Rapid HIT ID, which are fully integrated systems.

### STR genotyping using Nanopore sequencing

ONT sequencing of RPA singleplex product revealed that between 60.4 % (for FGA) and 96.3% (for TH01) of mapped reads were mapped to a true allele. This is somewhat lower than for Illumina sequencing in which over 94% of mapped reads were mapped to correct alleles. This reflects the lower general sequencing accuracy of Nanopore sequencing data, which is especially pronounced in the repetitive and homopolymeric regions characterizing STR loci^22^. **Supplementary Table S1** compares mapping percentages of ONT sequencing results generated in this study to mapping percentages of previously published PCR-based ONT sequencing of STR loci^9^. The comparison shows that the proportion of total reads assigned to a locus and reads mapped to an allele is considerably higher in our study compared to the 2020 study ^9^. In the same study, the genotypes generated by Illumina sequencing were all in concordance with the ones generated by the benchmarking method, while some loci were consistently genotyped incorrectly by ONT sequencing. Consistent failing of certain STR loci in ONT sequencing was reported in two other studies^8,10^ in which nanopore sequencing was performed on samples prepared using the commercially available Verogen ForenSeq multiplex kit (Verogen, San Diego, USA). These results, along with earlier studies^23^ point to the lack of maturity of Nanopore sequencing technology for forensic STR genotyping at the time. Since correct genotypes in our study were generated from a pooled singleplex, in contrast to the multiplex assay in the 2020 Tytgat paper, parallels cannot be drawn between the two amplification methods due to added complexity of the multiplex in the PCR assay. However, concordant genotyping of RPA singleplex in this study points to the improvement of ONT sequencing in dealing with repetitive regions. Recent advancements in both sequencing technology and data analysis pipelines^24^ are narrowing the performance gap between Illumina and ONT sequencing platforms for STR profiling. While Nanopore sequencing workflow is not considerably faster than Illumina sequencing or CE workflows, ONT’s MinION device comes with a low upfront cost of around 2000 USD, considerably lower than Illumina or CE setups. Reliable forensic genotyping ONT sequencing workflow is a potential solution for low-resource communities that are in regular or occasional need for forensic STR genotyping. Beside the cost, MinION device is portable and USB powered. Recent pilot study^25^ demonstrates fully of-grid forensic STR Nanopore sequencing in the field. If fully developed and validated, on-field Nanopore sequencing can become a valuable tool in the forensic genotyping toolbox, especially in specific situations such as disaster scenes in remote locations. Because of RPAs ability to reduce stutter^3^, its incorporation in ONT forensic sequencing workflow may lead to easier interpretation of the profiles resulting from ONT sequencing.

## Material and methods

### Sample collection and DNA extraction

In total, nine samples were utilized in this study. Of those, four DNA samples were extracted from blood of healthy volunteers, three DNA samples were extracted from buccal swabs and two were commercially available forensic controls. Ethical approval was granted by the ethical review board of Ghent University Hospital (approval No. BC-05557). All volunteers provided a signed informed consent prior to participation and all experiments were performed in accordance with relevant guidelines and regulations. The commercial controls were 9947A and 9948 forensic reference samples purchased from OriGene (Rockville, Maryland, USA). Eight samples were tested in the RPA followed by CE workflow. A subset of four DNA samples was used for the Illumina and Nanopore sequencing workflows; the two commercial controls and samples 3 and 4. Blood was collected via finger puncture using a 21G Minicollect® Lancelino safety lancet, with a penetration depth of 2.4 mm (Greiner Bio-One, Kremsmünster, Austria). The blood was collected into K3E K3EDTA Minicollect® collection tubes (Greiner Bio-One, Kremsmünster, Austria). DNA extraction from the collected blood samples was performed using the DNeasy® Blood and Tissue Kit (Qiagen, Hilden, Germany), according to the manufacturer’s instructions. DNA was extracted from buccal swabs using chelex resin purchased from Bio-Rad (Bio-Rad Laboratories, CA, USA) by the protocol outlined by Sweet et al.^26^. Extracted DNA was quantified using a Qubit 2.0 fluorometer (Thermo Fisher Scientific, Waltham, MA, USA).

### Benchmarking using capillary electrophoresis

The benchmark genotyping of the DNA extracted from the donor samples was conducted using PCR followed by amplicon sizing by capillary electrophoresis (CE). For each sample, 1 ng of DNA was amplified using the AmpFlSTR® Identifiler® Plus PCR Amplification Kit (Thermo Fisher Scientific, Waltham, MA, USA), according to the protocol provided by the manufacturer, opting for a 28 PCR cycles option. Resulting amplicons were analyzed using ABI3130xl Genetic Analyzer (Thermo Fisher Scientific, Waltham, MA, USA) equipped with 36 cm capillaries containing POP-4™ matrix (Applied Biosystems) according to the manufacturer’s instructions. 1 µl of PCR sample amplicons was mixed with 12 µl of Hi-Di™ formamide (Applied Biosystems) and 0.2 µL GeneScan™ 600 LIZ internal lane standard (Applied Biosystems). A run module with a 4 seconds injection time at 1.2 kV injection voltage and a separation time of 1550 seconds at 13 kV was used. The resulting electropherograms were interpreted using GeneMapper ID-x 1.2 software (Thermo Fisher Scientific, Waltham, MA, USA). Relative fluorescent units (RFU) analytical threshold was set to 100. For reference, the genotypes and source material of all analyzed samples are available in **Supplementary Tables S2 and S3**.

### RPA amplification, purification and pooling

Amplification of the samples was performed using the TwistAmp Liquid Basic Kit (TwistDx, Maidenhead, UK). The Reaction Buffer, 10x Basic E-mix, and 20x Core Reaction Mix were combined in the amounts required to achieve a 1X concentration in the final reaction mix. The concentration of dNTPs (Thermo Fisher Scientific, Waltham, MA, USA) in the reaction mix was 1.8 mM. The master mix was then added to the primer mixes, followed by the addition of 1 ng of DNA. The reaction was initiated by adding MgOAc at a reaction mix concentration of 14 mM. Based on their length, five loci (Amelogenin, D18S51, D21S11, D8S1179, TH01) were selected to be amplified using fluorescent primers (**Supplementary Table S4**), and analyzed by CE. Due to limited performance of the fiveplex, a panel of different loci was selected to be amplified in multiplex and analyzed by sequencing. The non-overlapping lengths of Amelogenin, D13S317, D16S539, D7S820, FGA, vWA allowed for the preliminary optimization by agarose gel screening (**Supplementary Figure S1**). Along with the sixplex, an expanded panel of thirteen loci was used: Amelogenin, D13S317, D16S539, D18S51, D21S11, D3S1358, D5S818, D7S820, D8S1179, FGA, TH01, TPOX, vWA. These were amplified in singleplex and thirteenplex. Since lengths of the thirteen loci overlap, assessment of the reaction success was not possible on the agarose gel. Widely used PCR primer sequences acquired from STRbase^27^ were adapted in length to match the recommended 30-35 nt (**Supplementary Table S4**). Sequences of the fluorescently labeled primers were the same for all loci except for D18S51 where two nucleotides were removed from the 5’ end in order to prevent the quenching of the fluorophore by the guanine residue. Due to the sensitivity of the assay to oligonucleotide concentrations in the master mix, the primer quantities varied across the 13 loci, ranging from 62,5 nM to 250 nM for each forward and reverse primer. The final primer concentration used in the singleplex reaction was determined by testing multiple concentrations and examining the product on a precast 2% agarose gel (Thermo Fisher Scientific, Waltham, MA, USA). The following concentrations were tested: 31,25 nM, 62,5 nM, 125 nM and 250 nM. For both positive and negative reactions, 3 µl of the amplification product was loaded on the gel. The concentration that gave the sharpest band and least visible smear on the agarose gel was selected for the experiment. The primer concentration ratios for the multiplex reactions were set based on their optimal concentration in the singleplex reaction and were adjusted not to exceed a maximal total oligo concentration of 2000 nM. Extensive optimization of multiplex primer concentrations was not conducted in this proof-of-concept study. Sequences and concentrations of individual primers can be found in **Supplementary Table S4**. Amplification was carried out for 40 minutes at 39°C. Post-amplification, the samples were purified using Zymo Clean and Concentrator-5 kits (Zymo Research, Irvine, CA, USA) according to the manufacturer’s instructions. Concentrations of the singleplex products were quantified using the PicoGreen assay (Thermo Fisher Scientific, Waltham, MA, USA), and subsequently pooled equimolarly. Since measuring of the DNA concentration of fluorescently labeled amplicons is not possible using Qubit or Picogreen techniques, pooling for CE analysis was performed by combining equal volumes of singleplex RPA product.

### Sensitivity assessment and mixture samples

Sensitivity assessment of singleplex RPA assay was conducted on serial dilutions of sample 2. DNA input amounts were: 500 pg, 250 pg, 125 pg, 62 pg and 31 pg. Amplification, purification and detection by CE was performed in the same manner as for 1 ng input samples.

### Illumina sequencing

The unlabeled RPA samples were sequenced on Illumina sequencing MiSeq platform, generally considered the golden standard for amplicon sequencing. The library preparation employed the PCR-free NEBNext® Ultra II for Illumina® kit (NEB, Ipswich, MA, USA), with input DNA ranging from 314 to 685 ng per sample. Following the ligation of TruSeq Unique Dual Index (UDI) adaptors, the libraries were purified using 0.9X AMPure XP beads (Beckman Coulter, High Wycombe, UK). Libraries were quantified using a Sequencing Library qPCR Quantification kit (Illumina, San Diego, CA, USA) and equimolarly pooled. Library quality was assessed using the Fragment Analyzer NGS Kit (Agilent Technologies, Santa Clara, CA, USA). The library pool was spiked with 25% PhiX. Paired-end 300 bp sequencing of the library pool was performed using a MiSeq Nano Flow Cell (Illumina, San Diego, CA, USA).

### Nanopore sequencing

Library preparation was conducted using the SQK-NBD114.24 kit, version 14, according to the manufacturer’s instructions (Oxford Nanopore Technology (ONT), Oxford, UK). For each library, 200 fmol of the purified RPA product was used, which corresponds to approximately 32 ng of DNA. Aside from reagents available in the ONT kit, additional reagents were used: NEB Next Ultra II End Repair/dA-Tailing Module was used for end preparation, the NEB Blunt/TA Ligase Master Mix for native barcode ligation, and the NEB Next Quick Ligation Module for adapter ligation (all from NEB, Ipswich, MA, USA). Quantification of the prepared libraries was performed using the Qubit dsDNA High Sensitivity Assay Kit on a Qubit fluorometer (Thermo Fisher, Waltham, MA, USA). A total of 12 distinct barcodes were used to create 12 separate libraries. For sequencing, 50 fmol of the pooled library was loaded onto a PromethION R10.4.1 flowcell, and the sequencing was conducted on a GridION device for a total duration of 40 hours.

## Data analysis

Nanopore sequencing reads were demultiplexed and filtered in real-time using Guppy basecaller (v7.2.13). Reads with a q score lower than 9 and shorter than 20 nt were filtered out. Reads passing the quality filter were later rebasecalled using the state-of-the-art Dorado basecaller (v0.5.1), with a high accuracy option. Paired-end Illumina reads were merged using the FLASH tool (v1.2.11) ^28^. Reads from both sequencing methods were processed using an in-house Python script pipeline. During the pipeline, amplicons are first extracted by matching to the RPA primer sequences using Fuzzy regex allowing two mismatches for nanopore data and zero mismatches for Illumina data. Extracted amplicons were then mapped using bwa mem (v0.7.17)^29^ to the library (**Supplementary File**) containing all known alleles present in >1% of the population, collected from STRbase 2.0^27^. Nanopore sequencing is inherently more noisy than Illumina sequencing. In order to reduce the noise, custom filtering based on the alignment score (AS), previously described by our group ^10^ was applied to the mapping data. Since the AS indicates the similarity between a read and its aligned reference, a higher AS typically indicates more accurate alignment to true alleles then to their +1 and -1 stutters. Conversely, reads affected by sequencing and basecalling errors tend to exhibit lower AS scores, potentially resulting in misalignment. The maximum achievable AS varies for each read and equals its span. The read span for each aligned read was determined based on the CIGAR string. All reads with an AS higher than 90% of the read span were retained for genotyping. No additional filtering was performed on Illumina data. Reads mapping to each allele were counted, and the allele with the highest count was called as present. If the allele with the second highest count was more than 50% of the highest count allele, it was also called as present ^10^. A different coverage threshold was set for each locus and it was calculated as 20 times the number of reference alleles for the given locus. If the coverage threshold was not met, genotyping was not performed.

## Conclusion

This study evaluates RPA as an amplification method within the standard STR typing workflows: CE and MPS. Sequencing using both Illumina and ONT technologies, as well as CE of the singleplex RPA product resulted in complete and accurate profiles across all samples and loci. By conducting a sensitivity assessment we demonstrated the successful amplification of four out of five analyzed loci from as little as 31 pg of template DNA. Dropouts were observed in locus D8S1179 for certain DNA inputs. This level of sensitivity is comparable to that typically achieved with optimized PCR-based STR amplification methods used in forensic DNA profiling. Illumina sequencing of the sixplex RPA product resulted in correct and complete STR profiles in all samples and loci except for one dropout in Amelogenin locus. These findings provide initial evidence for the feasibility of using RPA in multiplex STR amplification. However, the effort to simultaneously amplify all thirteen STR loci using RPA failed to produce complete and accurate profiles after DNA sequencing analysis. Similarly, CE analysis of a five-locus RPA multiplex failed to generate complete and accurate profiles. This outcome suggests that the current RPA chemistry may struggle with the increased complexity of a full forensic STR panel when amplified simultaneously. Even at a lower level of multiplexing, challenges remain in balancing amplification efficiency across multiple loci. Therefore, further optimization is required before RPA multiplex can be reliably used for generating multiplexed forensic STR profiles.

## Data availability

Sequencing data related to this article is available at the NCBI Sequence Read Archive using the BioProject ID: PRJNA1152328.

## Author contributions statement

**Sonja Škevin:** Conceptualization, Methodology, Investigation, Formal analysis, Visualization, Writing – original draft, Funding acquisition.

**Liesl De Keyzer:** Investigation, Formal analysis, Writing – review & editing.

**Lynn De Waele:** Investigation, Formal analysis, Writing – review & editing.

**David Van Hoofstat**: Investigation, Formal analysis.

**Olivier Tytgat:** Conceptualization, Investigation, Formal analysis, Writing – review & editing.

**Dieter Deforce:** Writing – review & editing; Supervision, Funding acquisition.

**Filip Van Nieuwerburgh:** Conceptualization, Methodology; Writing – review & editing; Supervision, Funding acquisition.

