## Supplementary File for "Recombinase Polymerase Amplification of Forensic Short Tandem Repeat Loci"

### Tables:

**Supplementary Table S1.** Comparison of sequencing metrics between Illumina and Nanopore sequencing data generated in this study and sequencing metric from the study by Tytgat and colleagues (14).

| Setup | ONT flowcell | Total number of reads | Reads assigned to a locus | Reads mapped To allele |
|---|---|---|---|---|
| Singleplex RPA (Illumina) | | 656774 | 65.41 % | 61.69 % |
| Singleplex RPA (ONT) | R10.4 | 8705545 | 63.60 % | 60.19 % |
| Multiplex PCR (Illumina) (14) | | 316450 | 36.60 % | 36.56 % |
| Multiplex PCR (ONT) (14) | R9.4 | 7263280 | 19.00 % | 16.34 % |
| | R10 | 2138545 | 18.86 % | 15.84 % |

**Supplementary Table S2.** True genotypes of the samples tested in this study as determined by capillary electrophoresis. Analysis used to test RPA amplicons is denoted in the row "Analysis", with MPS including both Illumina and ONT sequencing.

| **Sample** | **Sample 1** | **Sample 2** | **Sample 3** | **Sample 4** |
|---|---|---|---|---|
| **Source** | Control (9947A) | Control (9948) | Blood | Swab |
| **Analysis** | CE&MPS | CE&MPS | MPS | CE&MPS |
| AMEL | X,X | X,Y | X,X | X,X |
| D13S317 | 11,11 | 11,11 | 12,12 | 8,12 |
| D16S539 | 12,11 | 11,11 | 11,12 | 9,11 |
| D18S51 | 19,15 | 15,18 | 11,16 | 17,14 |
| D21S11 | 30,30 | 29,30 | 30.2,30 | 30,31 |
| D3S1358 | 14,15 | 15,17 | 16,16.1 | 17,17 |
| D5S818 | 11,11 | 13,11 | 13,11 | 11,12 |
| D7S820 | 10,11 | 11,11 | 10,9 | 12,9 |
| D8S1179 | 13,13 | 13,12 | 10,14 | 15,14 |
| FGA | 24,23 | 24,26 | 24,19 | 22,19 |
| TH01 | 9.3,8 | 6,9.3 | 6,9.3 | 8,8 |
| TPOX | 8,8 | 9,8 | 11,9 | 8,11 |
| vWA | 17,18 | 17,17 | 16,17 | 17,17 |

**Supplementary Table S3.** True genotypes of the samples tested in this study as determined by capillary electrophoresis. Analysis used to test RPA amplicons is denoted in the row "Analysis", with MPS including both Illumina and ONT sequencing.

| **Sample** | **Sample 5** | **Sample 6** | **Sample 7** | **Sample 8** | **Sample 9** |
|---|---|---|---|---|---|
| **Source** | Swab | Swab | Blood | Blood | Blood |
| **Analysis** | CE | CE | CE | CE | CE |
| AMEL | X,Y | X,X | X,X | X,X | X,X |
| D13S317 | 12,12 | 11,12 | 11,12 | 11,12 | 12,13 |
| D16S539 | 9,13 | 12,13 | 8,12 | 12,13 | 10,12 |
| D18S51 | 14,15 | 17,20 | 13,14 | 17,20 | 14,16 |
| D21S11 | 28,30. | 30,33.2 | 32,32.2 | 30,33.2 | 29,29 |
| D3S1358 | 16,17 | 16,16 | 16,17 | 16,16 | 14,18 |
| D5S818 | 12,12 | 10,11 | 11,12 | 10,11 | 11,12 |
| D7S820 | 9,10 | 7,10 | 11,12 | 7,10 | 8,11 |
| D8S1179 | 14,16 | 13,15 | 11,14 | 13,15 | 13,15 |
| FGA | 21,24 | 22,24 | 20,24 | 22,24 | 20,22 |
| TH01 | 8,9.3 | 6,9.3 | 9,9.3 | 6,9.3 | 9,9.3 |
| TPOX | 9,11 | 8,11 | 8,8 | 8,11 | 10,11 |
| vWA | 15,19 | 16,16 | 17,18 | 16,16 | 16,17 |

**Supplementary Table S4.** Sequences and concentrations of labeled primers used in RPA followed by capillary electrophoresis. Fluorophores are denoted in bold letters at the 5' end of the primer sequence.

| Locus | Direction | $c_{in\ singleplex}$ (nM) | $c_{in\ multiplex}$ (nM) | Primer sequence |
|---|---|---|---|---|
| Amelogenin | Forward | 125 | 62,5 | **/VIC/-**CGTTAACAATGCCCTGGGCTCTGTAAAGAA |
| | Reverse | 125 | 62,5 | CCAACCATCAGAGCTTAAACTGGGAAGCTG |
| D8S1179 | Forward | 250 | 125 | ATTGCAACTTATATGTATTTTTGTATTTCAT |
| | Reverse | 250 | 125 | **/VIC/**-ACCAAATTGTGTTCATGAGTATAGTTTC |
| D18S51 | Forward | 125 | 62,5 | **/56-FAM/**CAGGAGGAGTTCTTGAGCCCAGAAGGTTA |
| | Reverse | 125 | 62,5 | ACCCGACTACCAGCAACAACACAAATAAAC |
| D21S11 | Forward | 125 | 62,5 | CTCCATAAATATGTGAGTCAATTCCCCAAG |
| | Reverse | 125 | 62,5 | **/56-FAM/**ATGTTGTATTAGTCAATGTTCTCCAGAGAC |
| TH01 | Forward | 125 | 62,5 | **/56-FAM/**TATCTGGGCTCTGGGGTGATTCCCATTGGCCTGTTC |
| | Reverse | 125 | 62,5 | GCACCGAAGACCCCTCCTGTGGGCTGAAAAGCTC |

**Supplementary table S5.** Unlabeled primers used in RPA assay followed by Illumina and ONT sequencing. For reference, lengths of longest alleles present in at least 1% of European population (http://spsmart.cesga.es/popstr.php) are included in the table.

| Locus | Longest allele | Length (bp) | Direction | Concentration in singleplex (nM) | Concentration in multiplex (nM) | Primer sequence |
|---|---|---|---|---|---|---|
| Amelogenin | Y | 129 | Forward | 250 | 62,5 | CGTTAACAATGCCCTGGGCTCTGTAAAGAA |
| | | | Reverse | | | CCAACCATCAGAGCTTAAACTGGGAAGCTG |
| D13S317 | 14 | 218 | Forward | 250 | 62,5 | TGGTATCACAGAAGTCTGGGATGTGGAGGA |
| | | | Reverse | | | GTTGAGCCATAGGCAGCCCAAAAAGACAGA |
| D16S539 | 14 | 300 | Forward | 250 | 125 | GGGGGTCTAAGAGCTTGTAAAAAG |
| | | | Reverse | | | ACCCGACTACCAGCAACAACACAAATAAAC |
| D18S51 | 21 | 342 | Forward | 250 | 31,3 | GGCAGGAGGAGTTCTTGAGCCCAGAAGGTTA |
| | | | Reverse | | | ACCCGACTACCAGCAACAACACAAATAAAC |
| D21S11 | 33.2 | 253 | Forward | 250 | 62,5 | CTCCATAAATATGTGAGTCAATTCCCCAAG |
| | | | Reverse | | | ATGTTGTATTAGTCAATGTTCTCCAGAGAC |
| D3S1358 | 19 | 173 | Forward | 125 | 31,3 | TATGTGACAAGGGTGATTTTCCTCTTTGGTATCC |
| | | | Reverse | | | TCCAATCATAGCCACAGTTTACAACATTTGTATCT |
| D5S818 | 14 | 167 | Forward | 62,5 | 31,3 | GAGCTATGATTCCCCCACTGCAGTCCAATCTGGGT |
| | | | Reverse | | | CATCTCTTATACTCATGAAATCAACAGAGGCTTGC |
| D7S820 | 13 | 259 | Forward | 125 | 62,5 | GGTTTCACCATGTTGGTCAGGCTGACTATG |
| | | | Reverse | | | ATAACGATTCCACATTTATCCTCATTGAC |
| D8S1179 | 16 | 242 | Forward | 125 | 31,3 | ATTGCAACTTATATGTATTTTTGTATTTCATG |
| | | | Reverse | | | ACCAAATTGTGTTCATGAGTATAGTTTC |
| FGA | 21 | 381 | Forward | 125 | 62,5 | GGCAGGAGGAGTTCTTGAGCCCAGAAGGTTA |
| | | | Reverse | | | ACCCGACTACCAGCAACAACACAAATAAAC |
| TH01 | 9.3 | 206 | Forward | 125 | 31,3 | TATCTGGGCTCTGGGGTGATTCCCATTGGCCTGTTC |
| | | | Reverse | | | GCACCGAAGACCCCTCCTGTGGGCTGAAAAGCTC |
| TPOX | 12 | 308 | Forward | 125 | 125 | CCAGAACCGTCGACTGGCACAGAACAGGCACTTAG G |
| | | | Reverse | | | TCGTGTTTGCGTCCCCAACGCTCAAACGTGAGGTTG |
| vWA | 20 | 163 | Forward | 125 | 31,3 | GCCCTAGTGGATGATAAGAATAATCAGTATGTG |
| | | | Reverse | | | GGACAGATGATAAATACATAGGATGGATGG |

**Figures:**

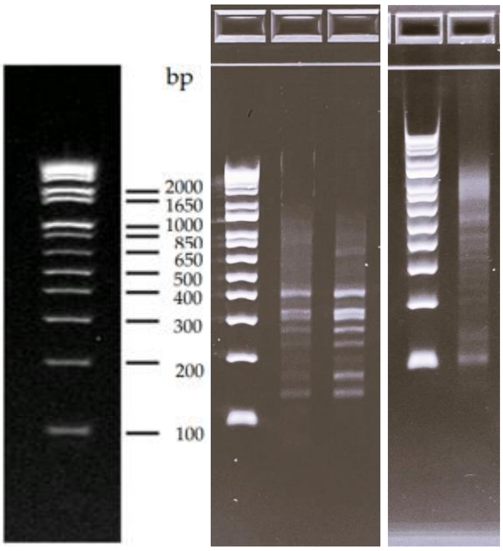

**Supplementary Figure S1**. Gel electrophoresis of sixplex and and thirteenplex RPA product on a 2 %agarose gel. Sizes of marker fragments are included in the leftmost side of the picture.

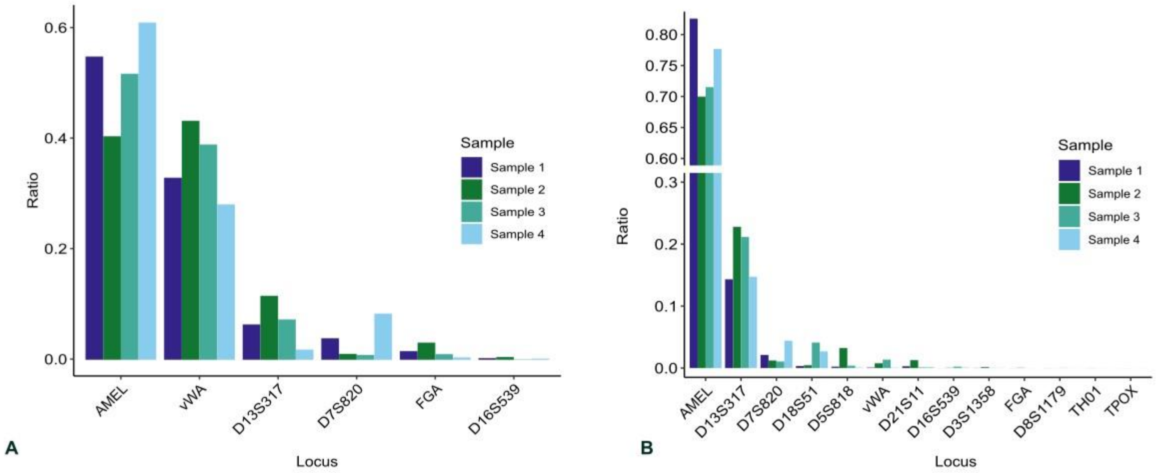

**Supplementary Figure S2**. **Amplification bias in RPA multiplexes, as measured by ONT sequencing.** The Y-axis shows the ratio of reads assigned to a certain locus divided by the sum of all reads assigned to any locus in **a.** Sixplex setup and **b.** Thirteenplex setup.

**Raw photographs of gel electrophoresis results:**

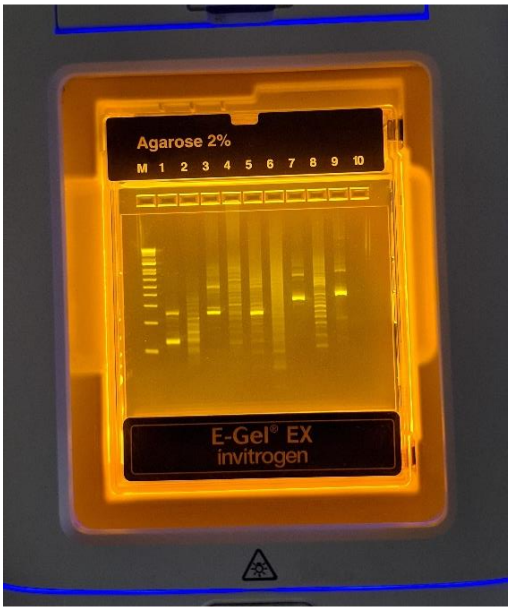

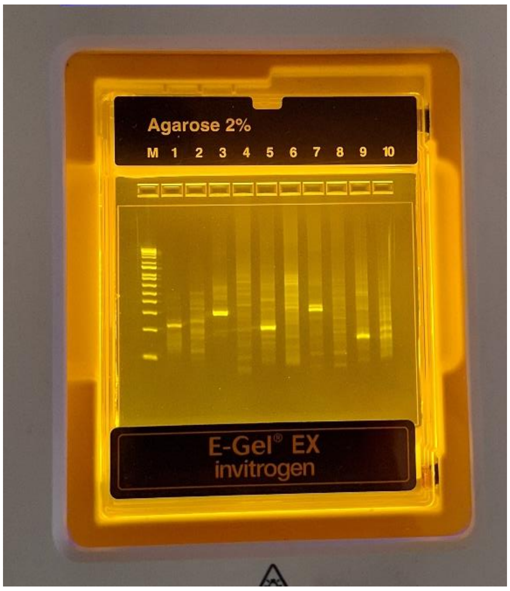

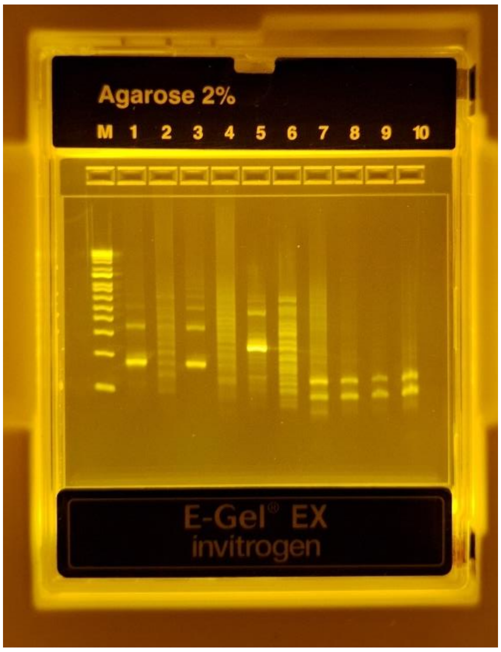

Applied Biosystems
GeneMapper® ID-X 1.2

| Sample Name | Panel | SQO | CGQ |
|---|---|---|---|
| NTC | | ✓ | ● |

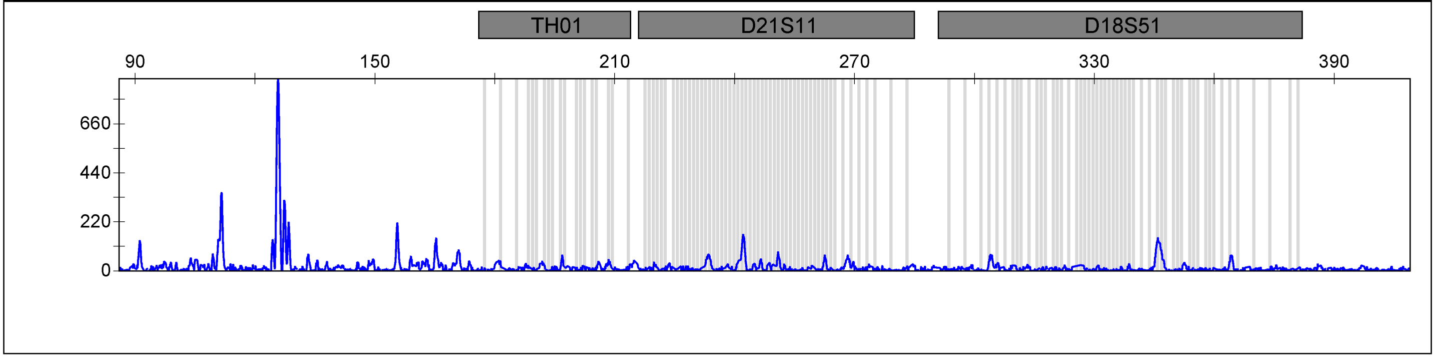

| | | | |
|---|---|---|---|
| | | ✓ | ● |

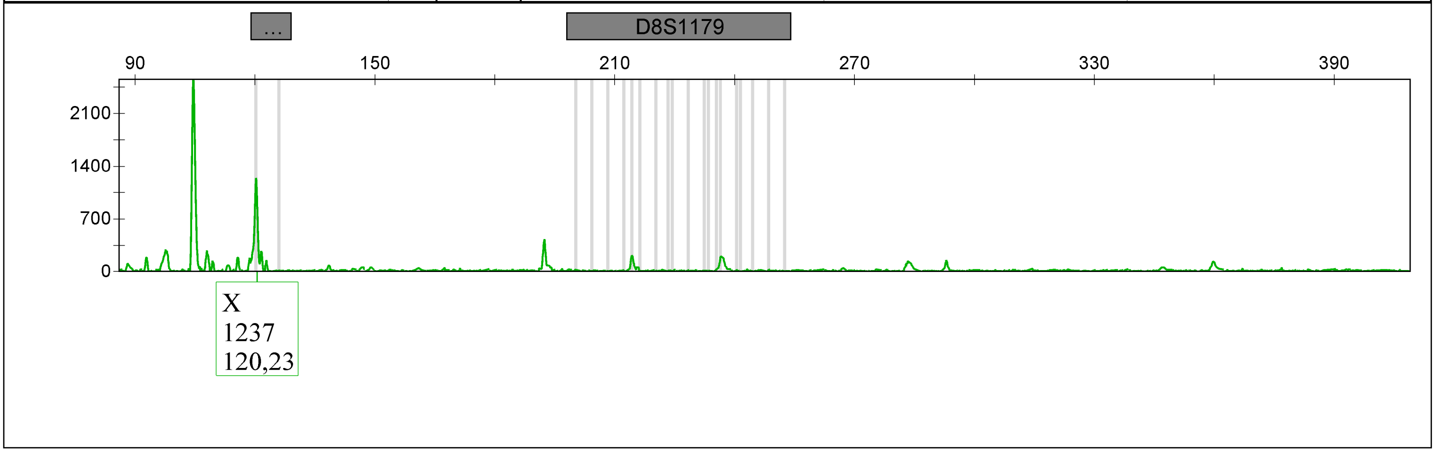

Applied Biosystems
GeneMapper® ID-X 1.2

Project: Sonja-3500-24-04-05

| Sample Name | Panel | SQO | CGQ |
|---|---|---|---|
| Sample 1 | RPA_panel | ✓ | ● |

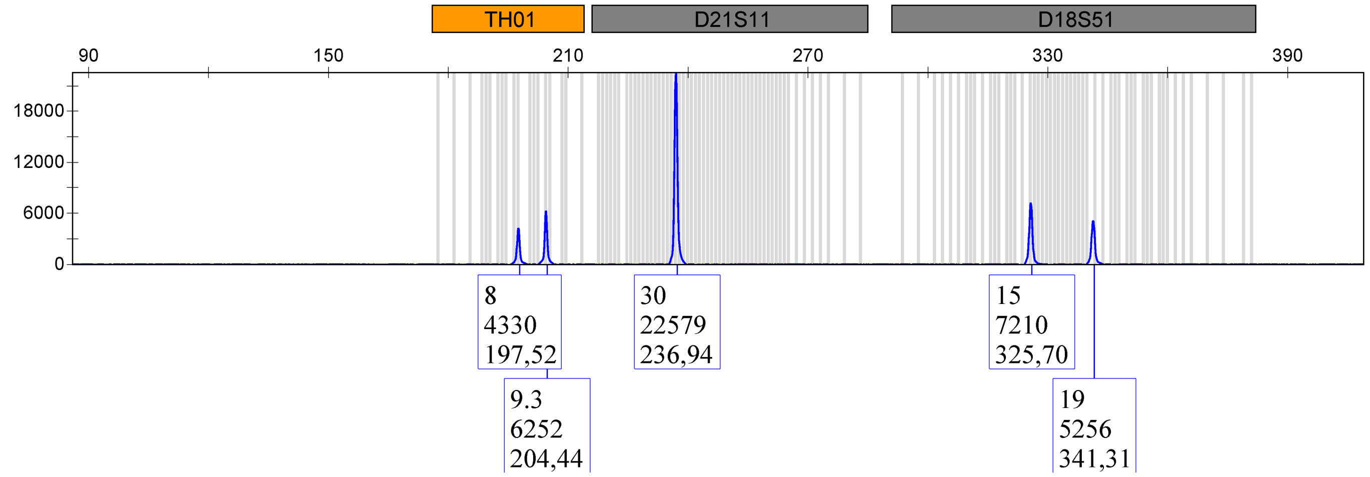

| | RPA_panel | ✓ | ● |
|---|---|---|---|

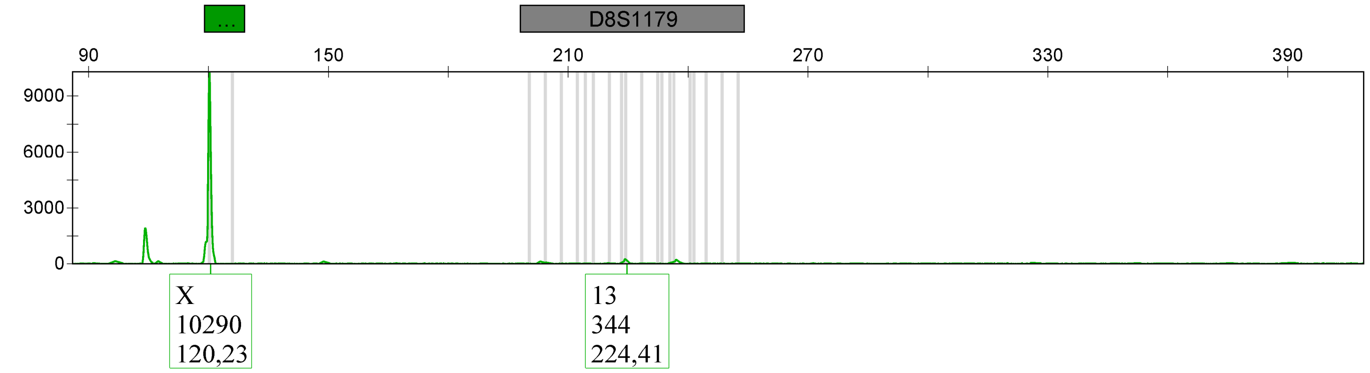

| Sample Name | Panel | SQO | CGQ |
|---|---|---|---|
| Sample 2 | RPA_panel | ✓ | |

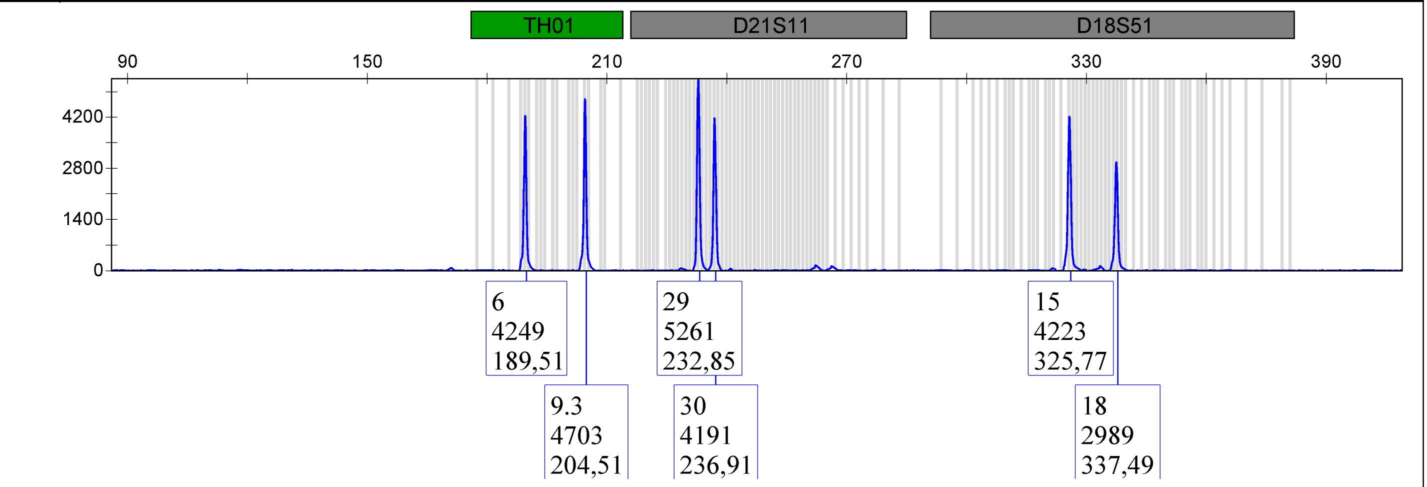

| | RPA_panel | ✓ |
|---|---|---|

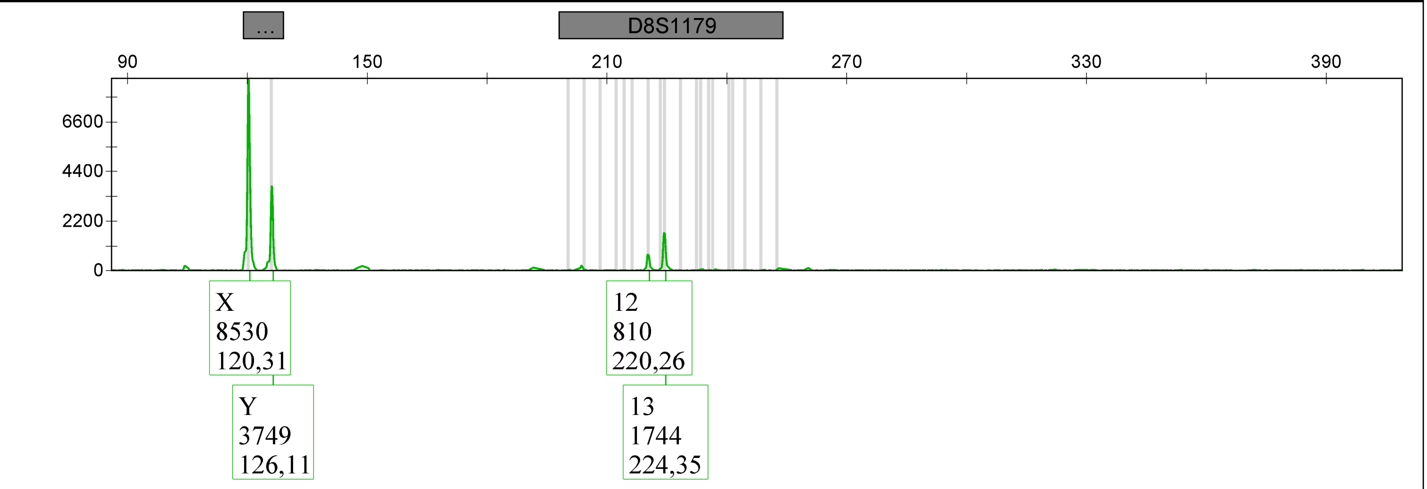

| Sample Name | Panel | SQO | CGQ |
|---|---|---|---|
| Sample 4 | RPA_panel | ✓ | |

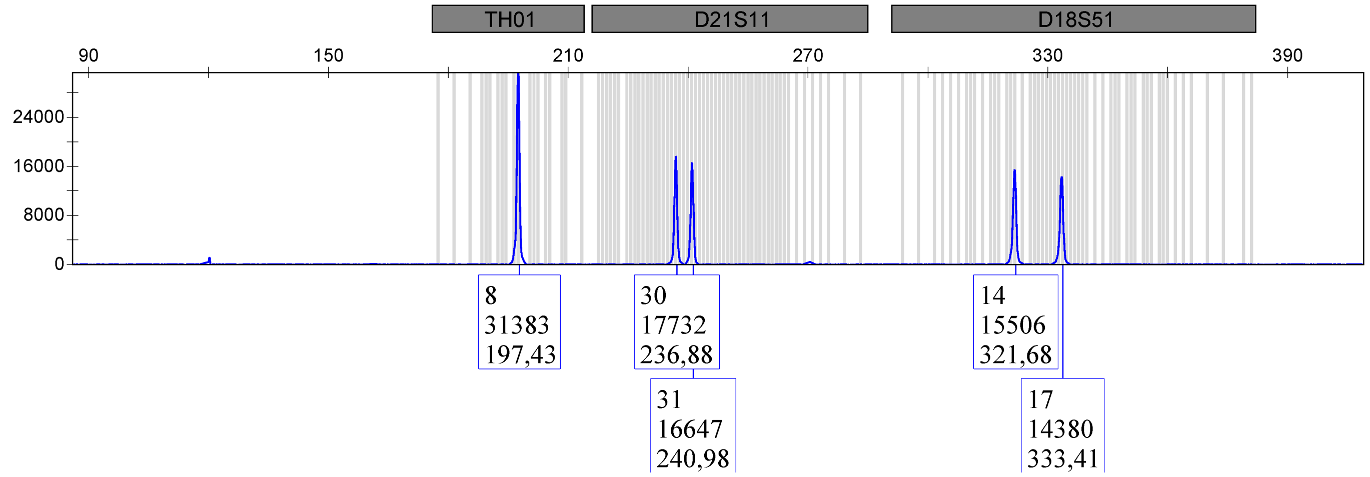

| | Panel | SQO | CGQ |
|---|---|---|---|
| | RPA_panel | ✓ | |

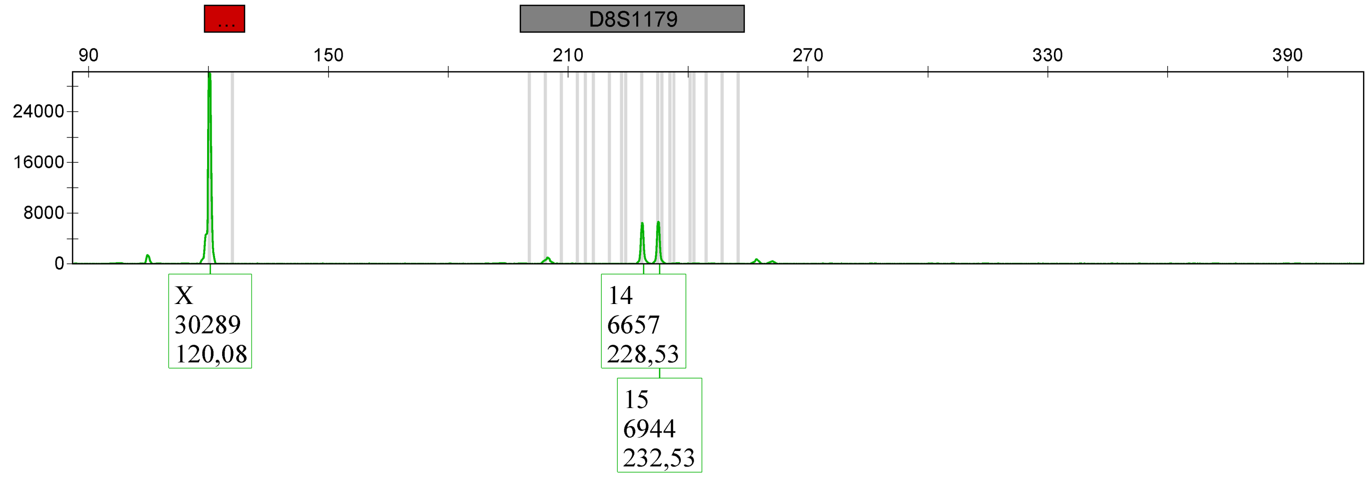

| Sample Name | Panel | SQO | CGQ |
|---|---|---|---|
| Sample 5 | RPA_panel | ✓ | ● |

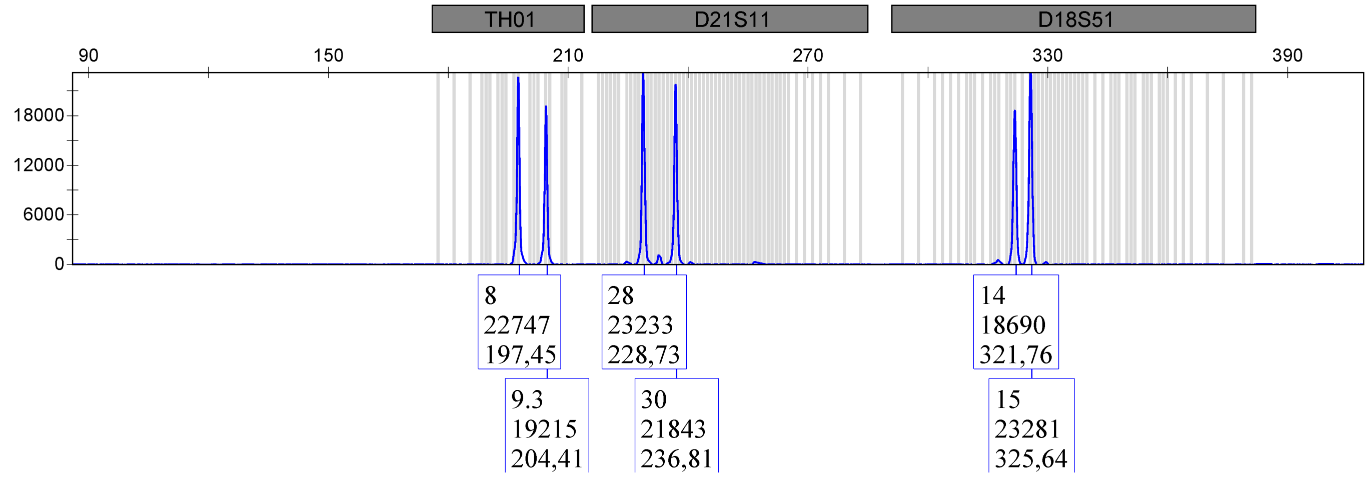

| | Panel | | |
|---|---|---|---|
| | RPA_panel | ✓ | ● |

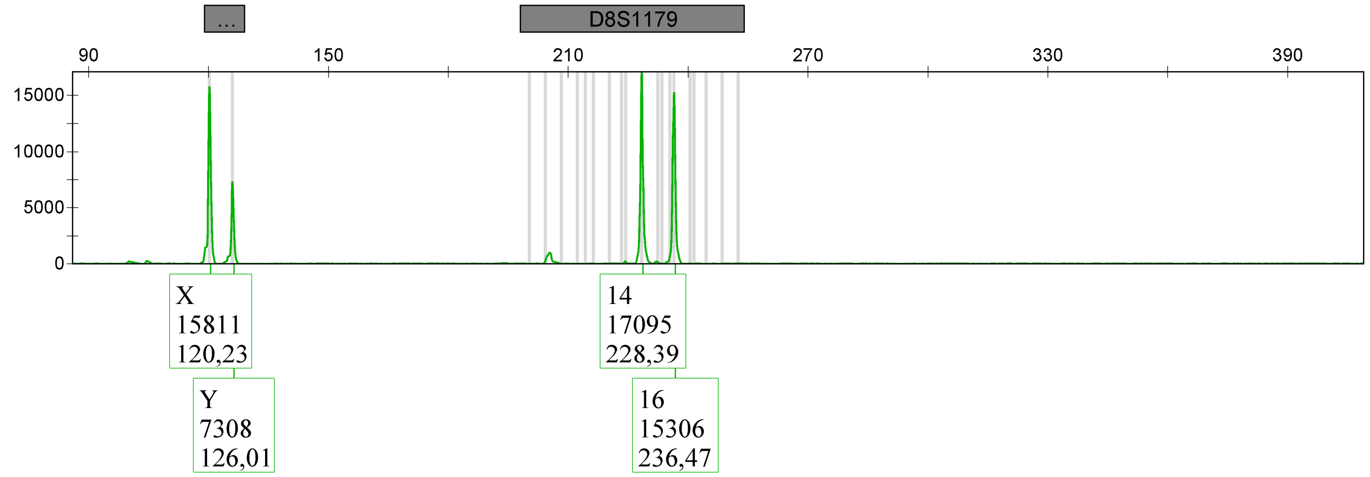

Applied Biosystems
GeneMapper® ID-X 1.2

Project: Sonja-3500-24-04-05

| Sample Name | Panel | SQO | CGQ |
|---|---|---|---|
| Sample 6 | RPA_panel | ✓ | |

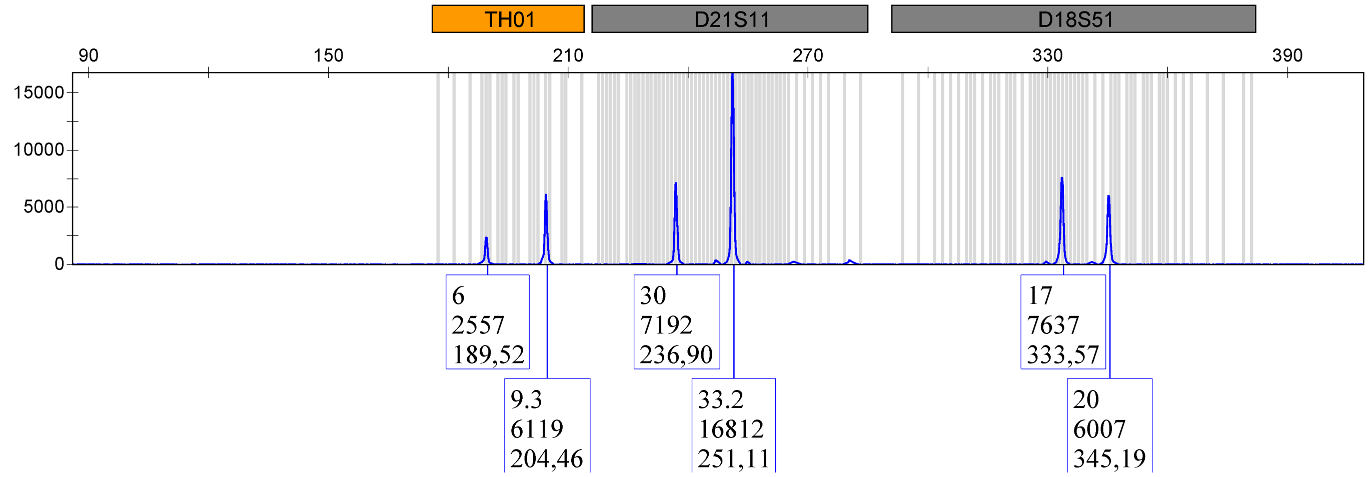

| | RPA_panel | ✓ |
|---|---|---|

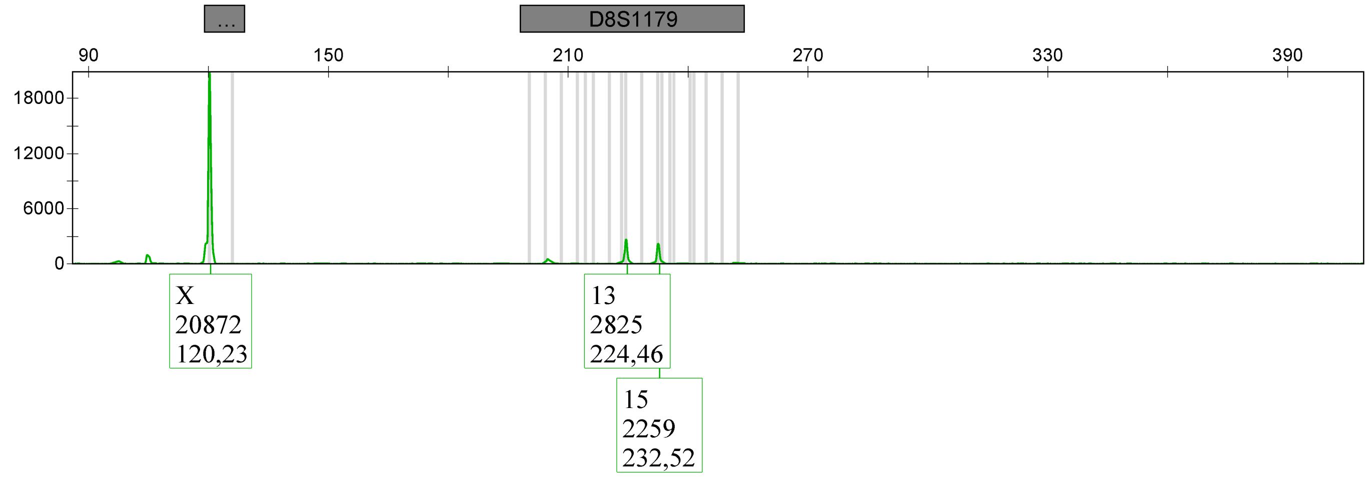

| Sample Name | Panel | SQO | CGQ |
|---|---|---|---|
| Sample 7 | RPA_panel | ✓ | |

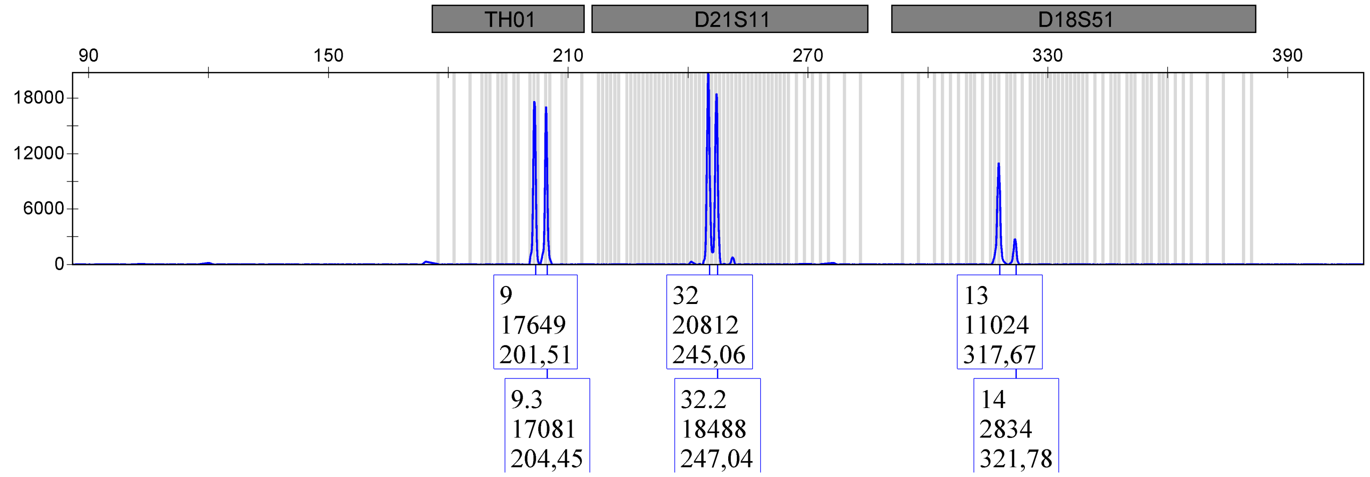

| | RPA_panel | ✓ |
|---|---|---|

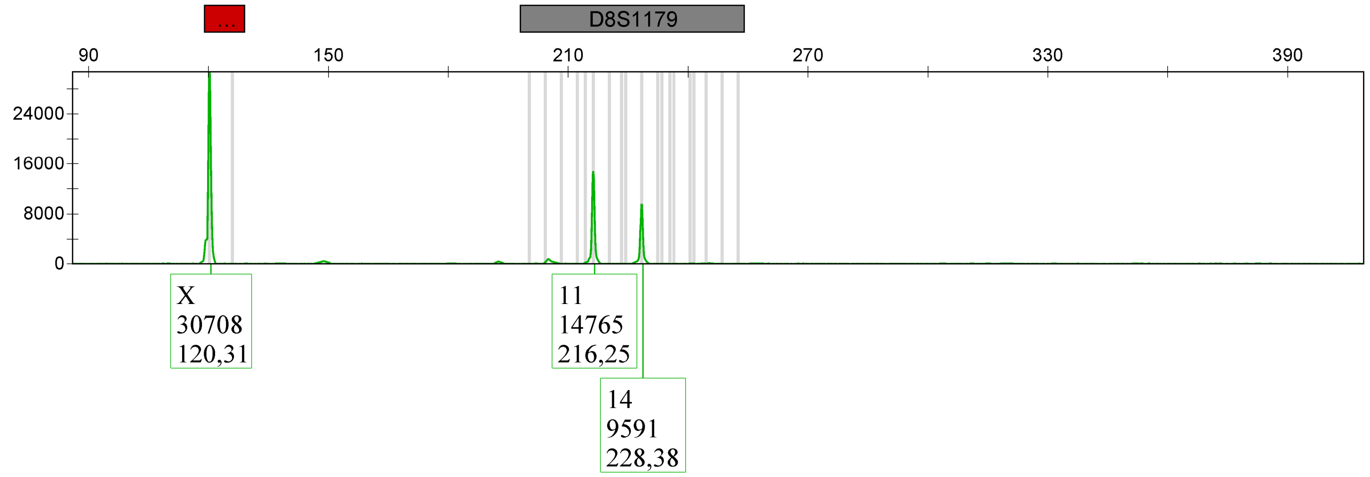

| Sample Name | Panel | SQO | CGQ |
|---|---|---|---|
| Sample 8 | RPA_panel | ✓ | |

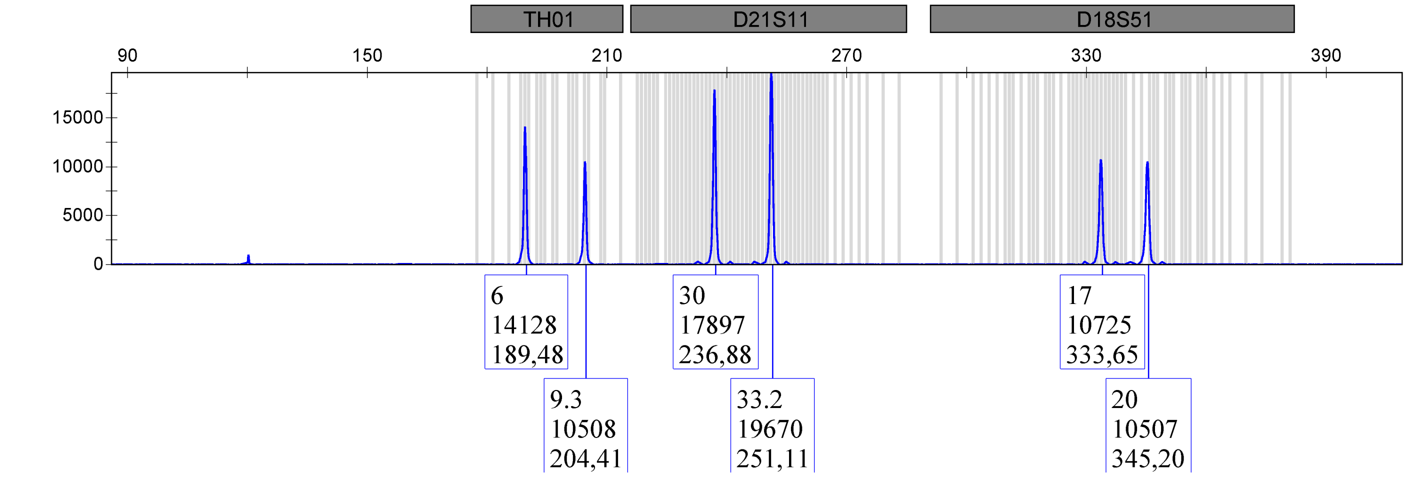

| | Panel | |
|---|---|---|
| | RPA_panel | ✓ |

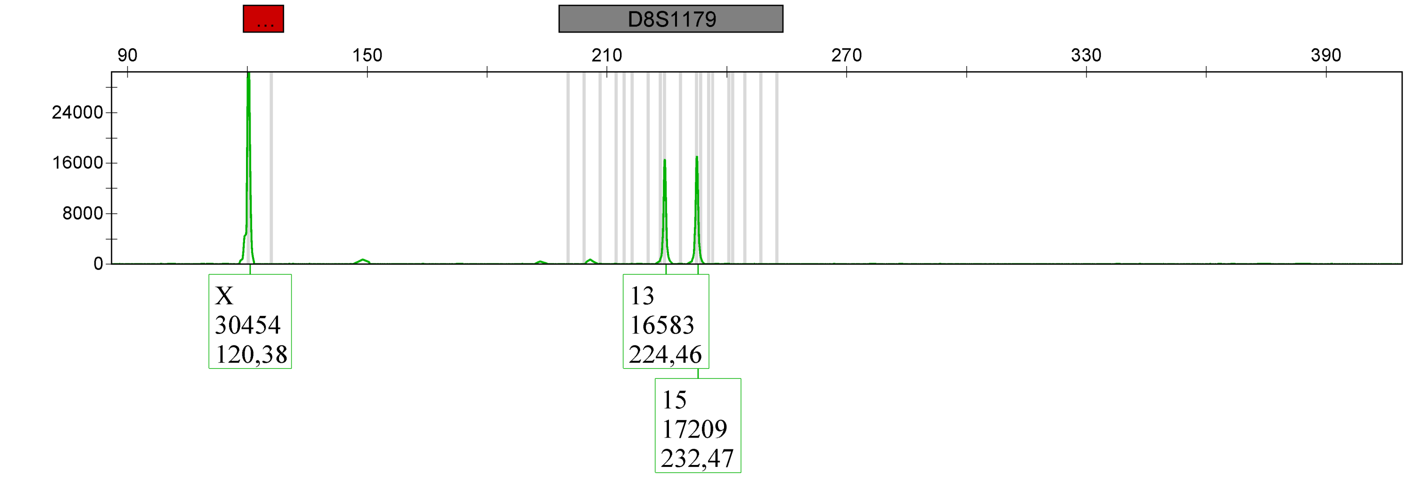

Project: Sonja-3500-24-04-05

| Sample Name | Panel | SQO | CGQ |
|---|---|---|---|
| Sample 9 | RPA_panel | ✓ | |

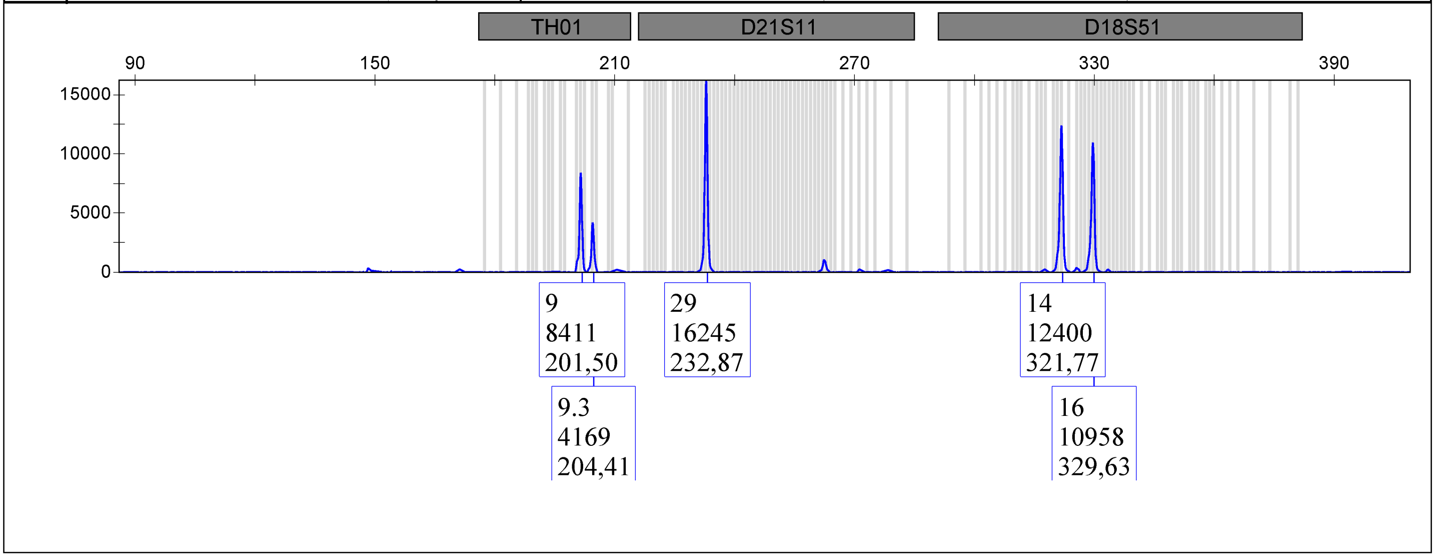

| Sample Name | Panel | SQO | CGQ |
|---|---|---|---|
| | RPA_panel | ✓ | |

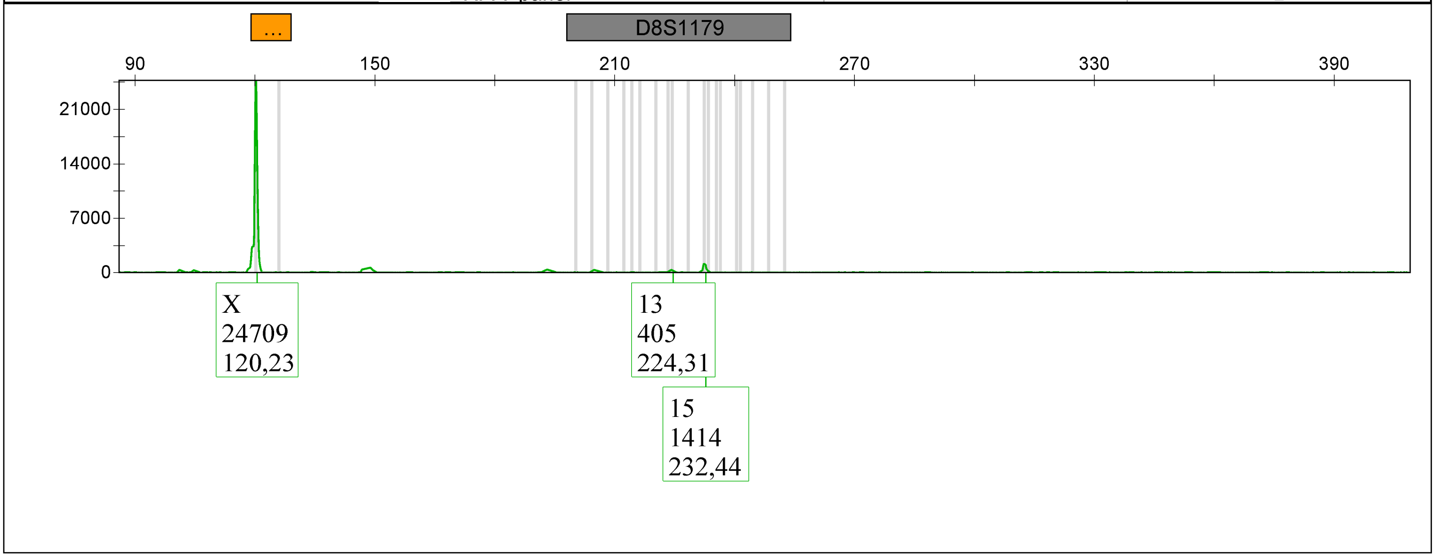

Applied Biosystems
GeneMapper® ID-X 1.2

| Sample Name | Panel | SQO | CGQ |
|---|---|---|---|
| NTC | RPA_panel | | |

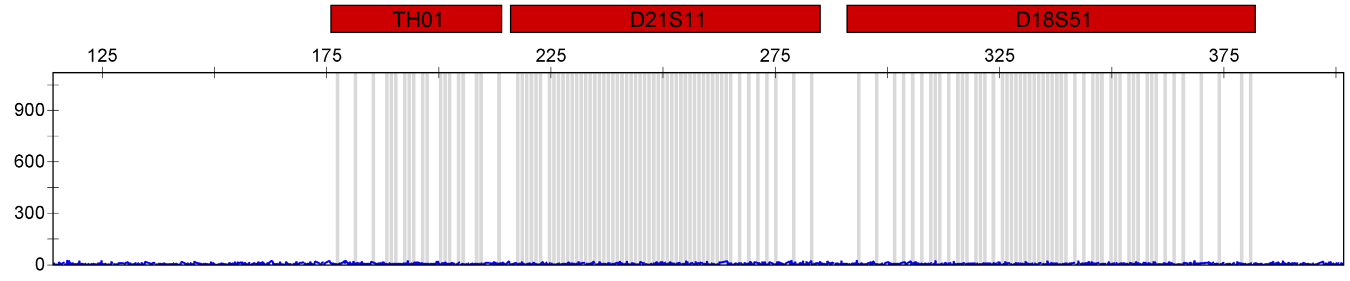

| | RPA_panel |
|---|---|

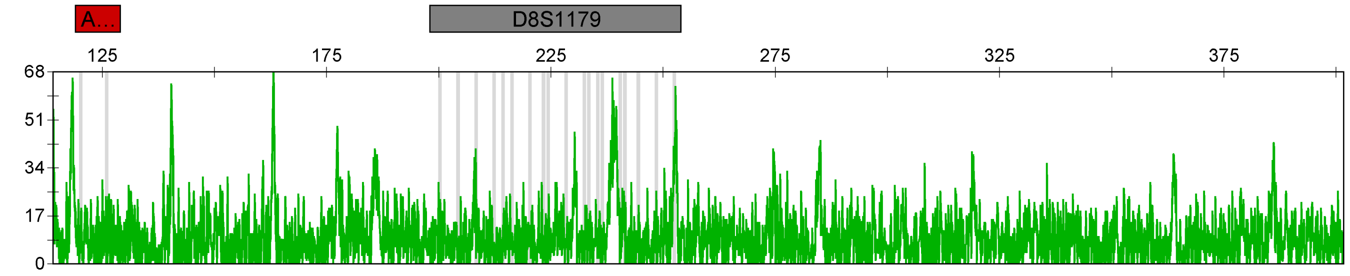

| Sample Name | Panel | SQO | CGQ |
|---|---|---|---|
| Sample 1 | RPA_panel | ✓ | |

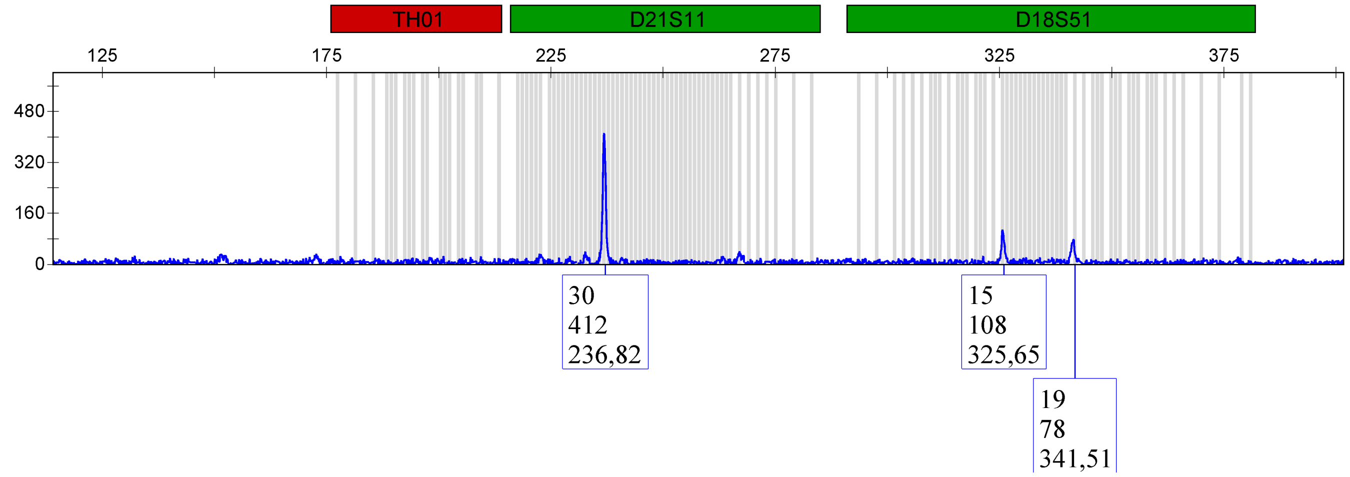

| | RPA_panel | ✓ |
|---|---|---|

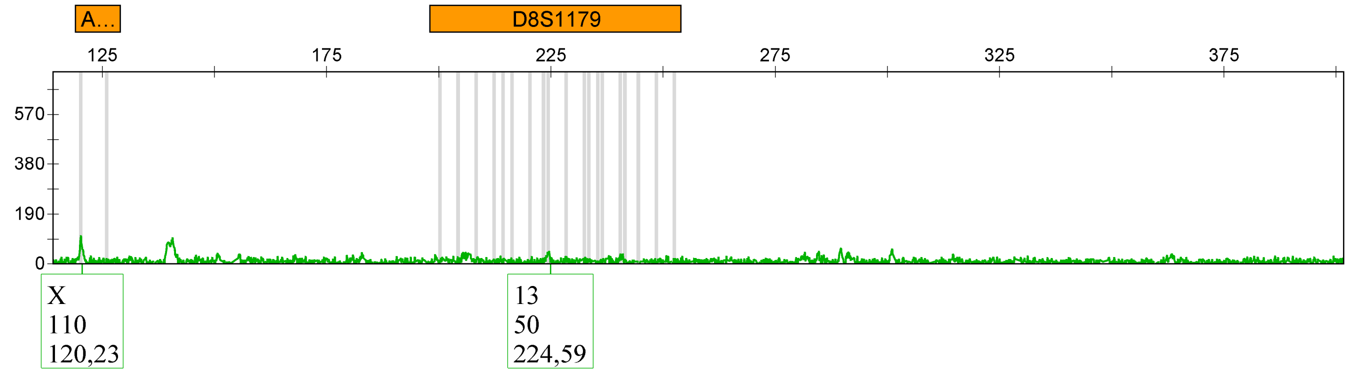

| Sample Name | Panel | SQO | CGQ |
|---|---|---|---|
| Sample 2 | RPA_panel | ✓ | |

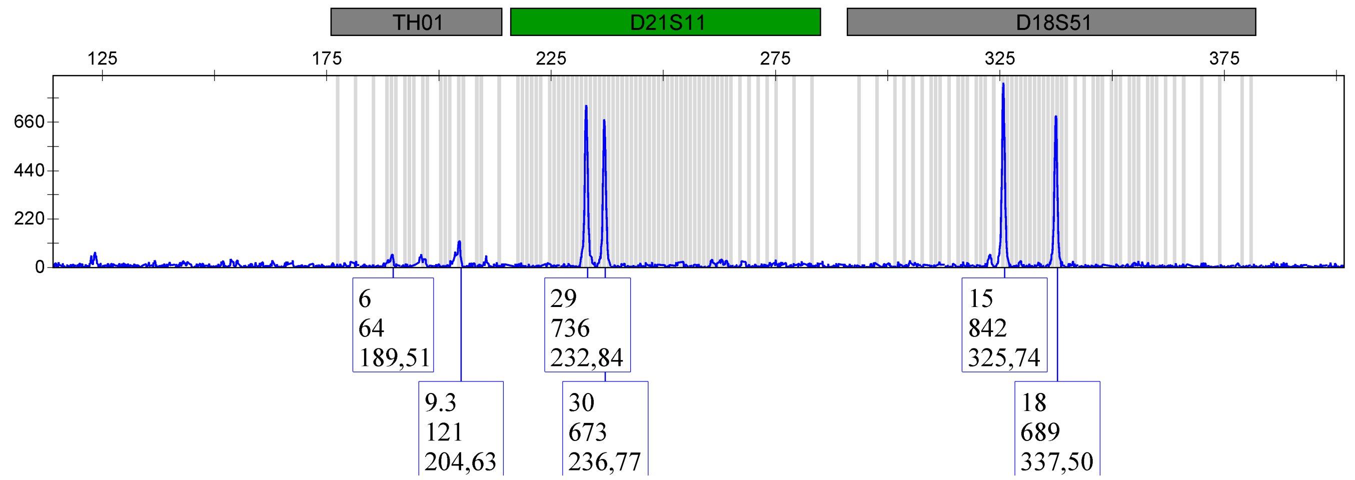

| | RPA_panel | ✓ |
|---|---|---|

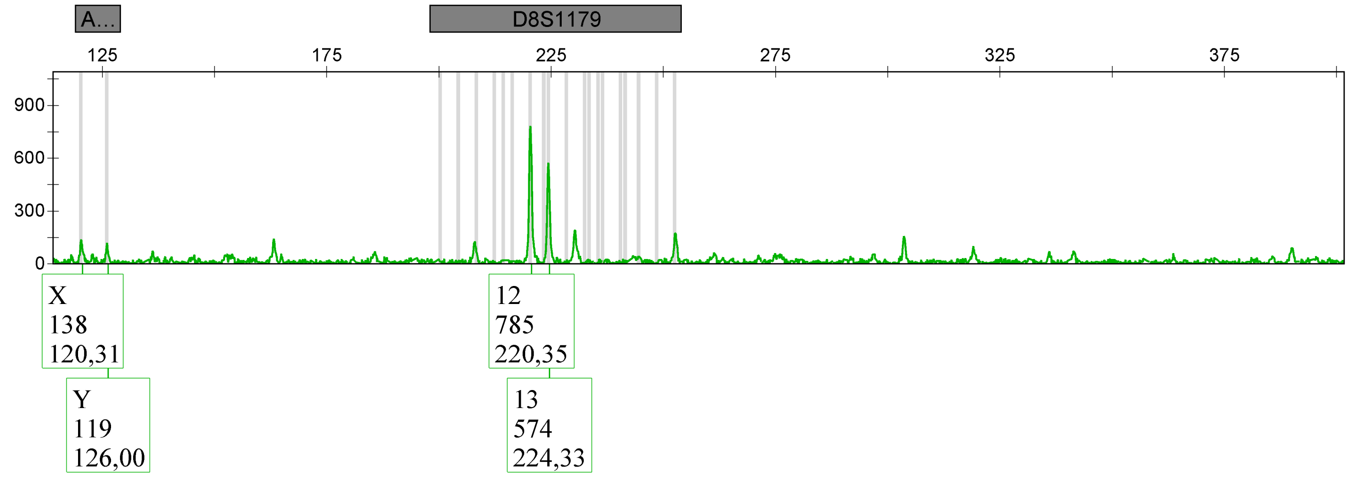

| Sample Name | Panel | SQO | CGQ |
|---|---|---|---|
| Sample 4 | RPA_panel | ✓ | |

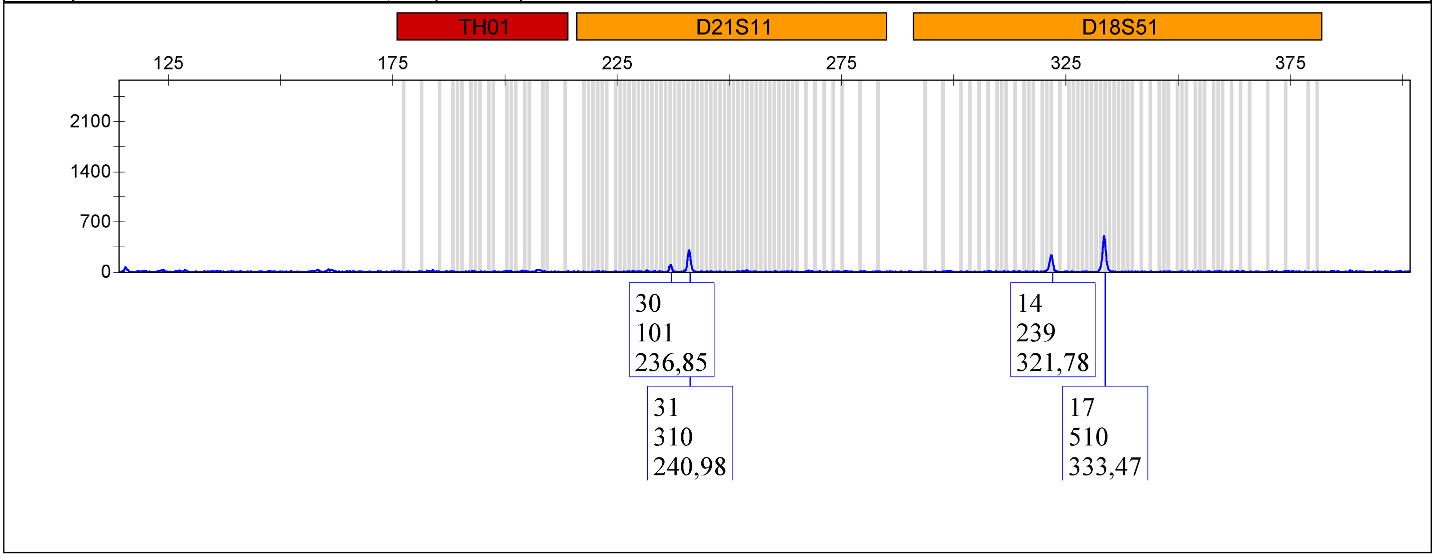

| | | |
|---|---|---|
| | [redacted] RPA_panel | ✓ |

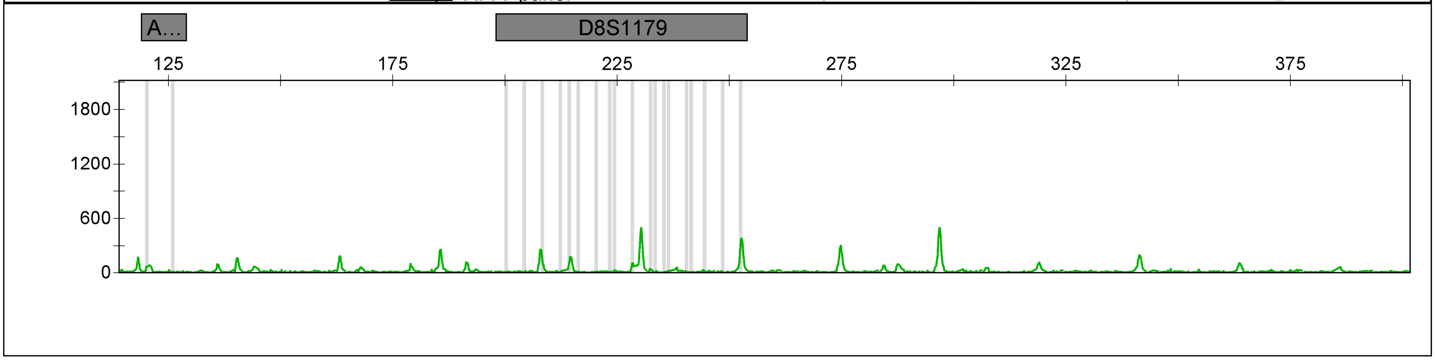

| Sample Name | Panel | SQO | CGQ |
|---|---|---|---|
| Sample 5 | RPA_panel | | |

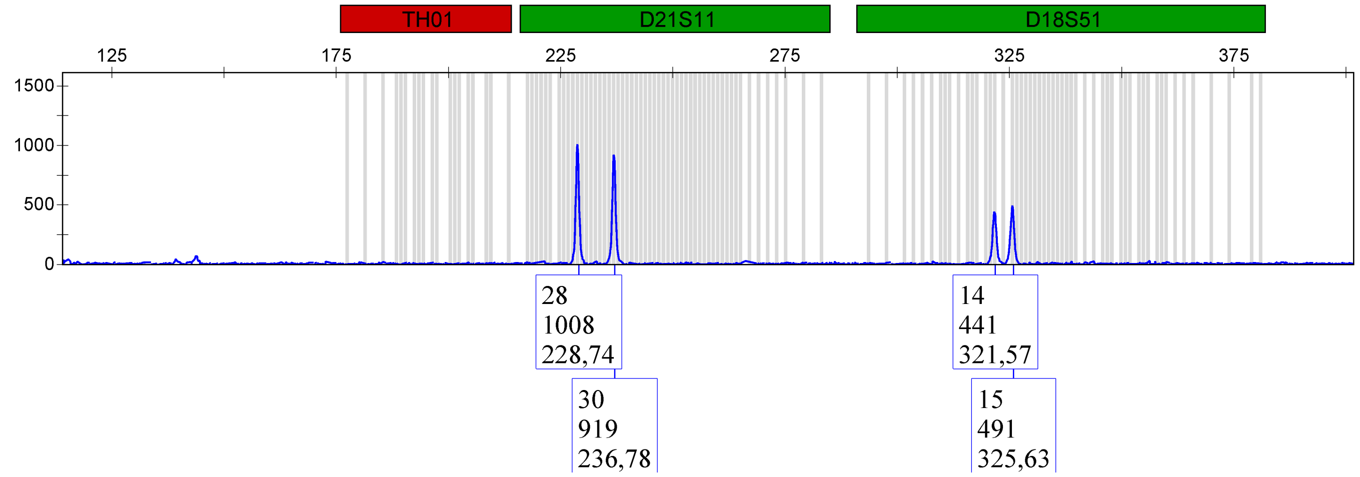

RPA_panel

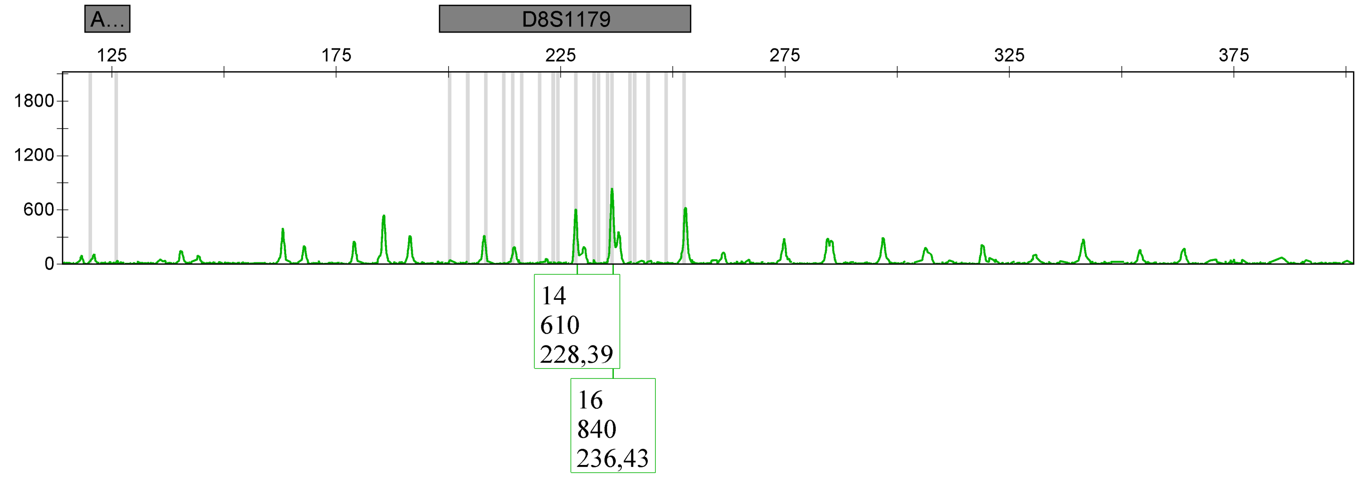

Applied Biosystems

GeneMapper® ID-X 1.2

| Sample Name | Panel | SQO | CGQ |
|---|---|---|---|
| Sample 6 | RPA_panel | ✓ | ● |

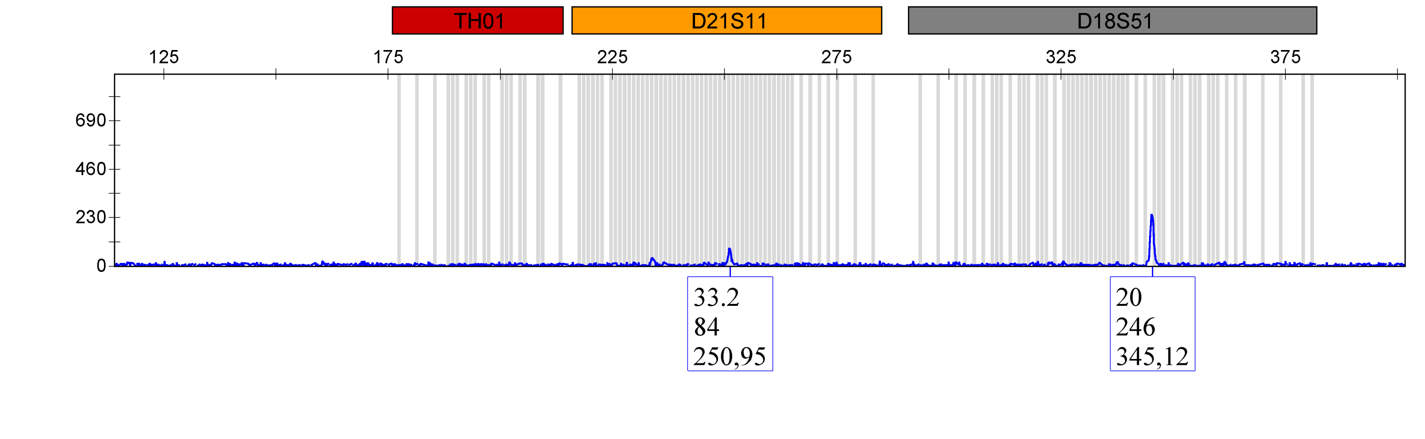

| | RPA_panel | ✓ | ● |
|---|---|---|---|

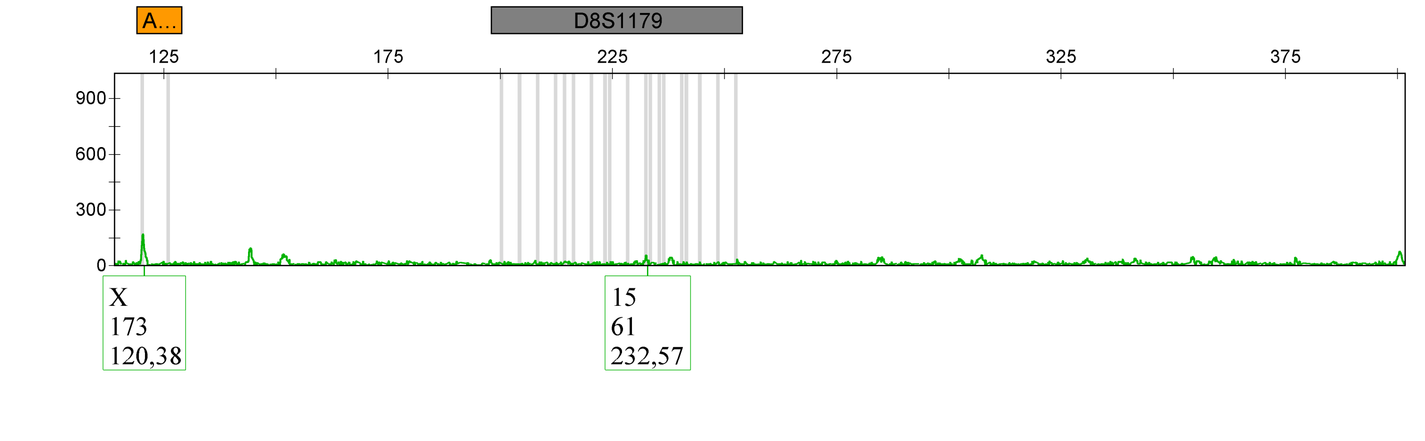

| Sample Name | Panel | SQO | CGQ |
|---|---|---|---|
| Sample 7 | RPA_panel | | |

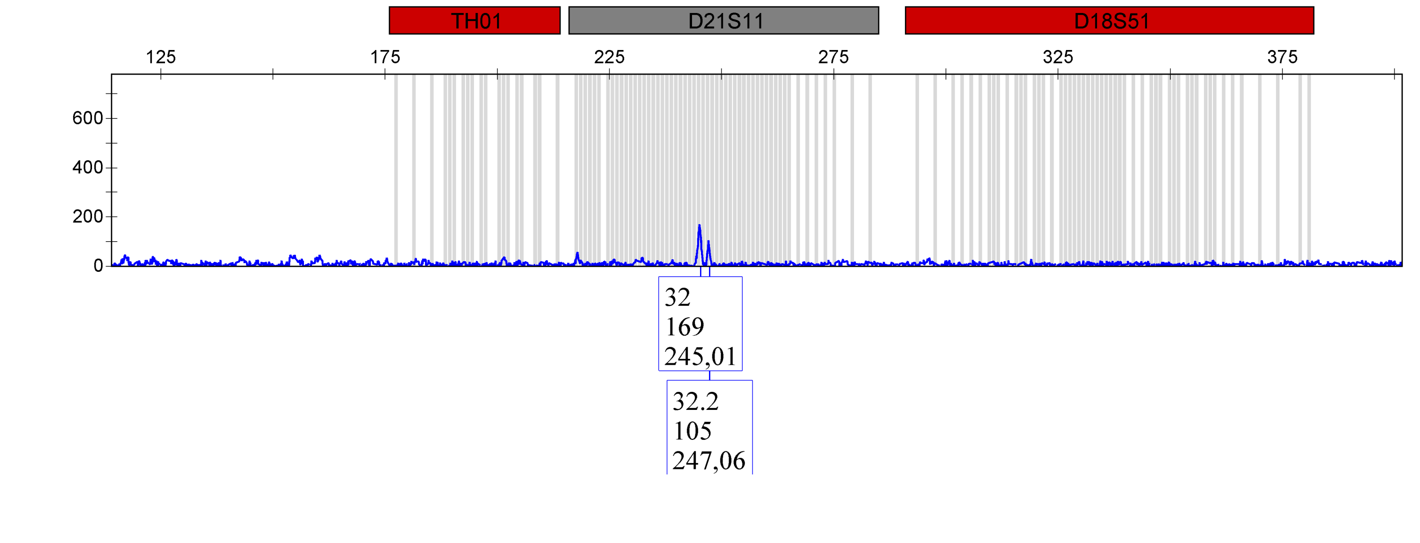

| | |
|---|---|
| | [redacted] RPA_panel |

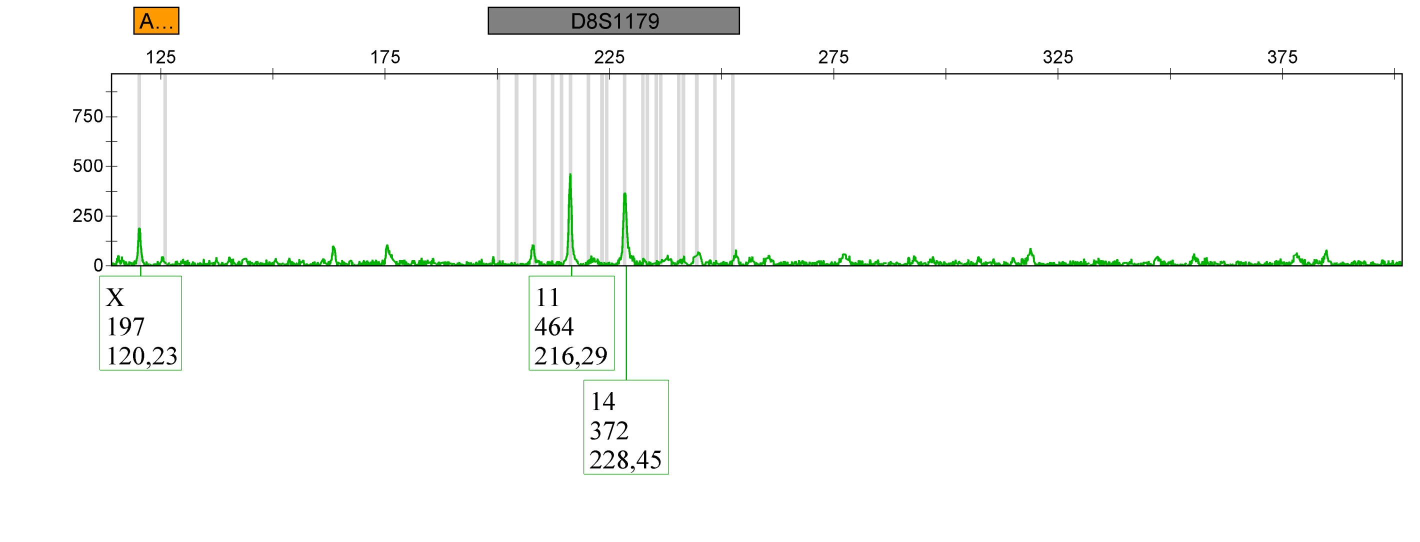

Applied Biosystems
GeneMapper® ID-X 1.2

| Sample Name | Panel | SQO | CGQ |
|---|---|---|---|
| Sample 8 | RPA_panel | ✓ | |

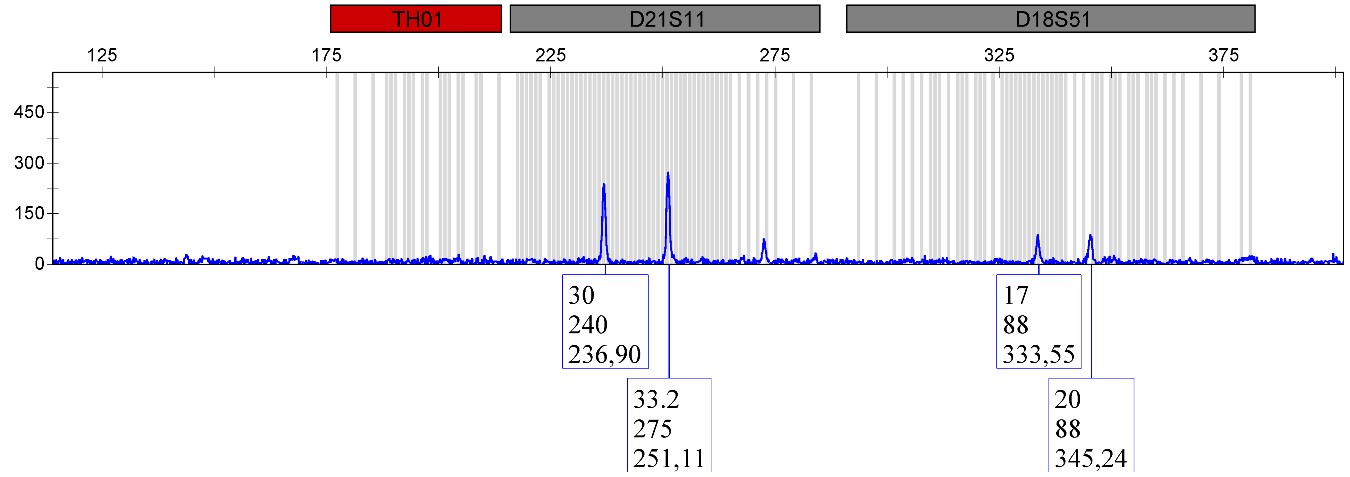

RPA_panel ✓

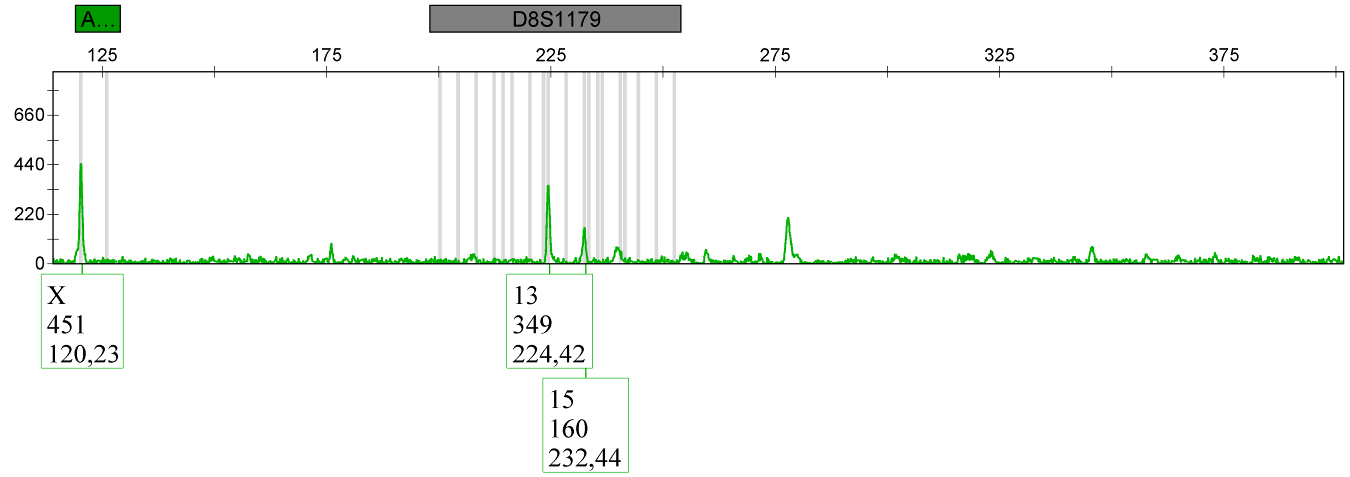

| Sample Name | Panel | SQO | CGQ |
|---|---|---|---|
| Sample 9 | RPA_panel | ✓ | ● |

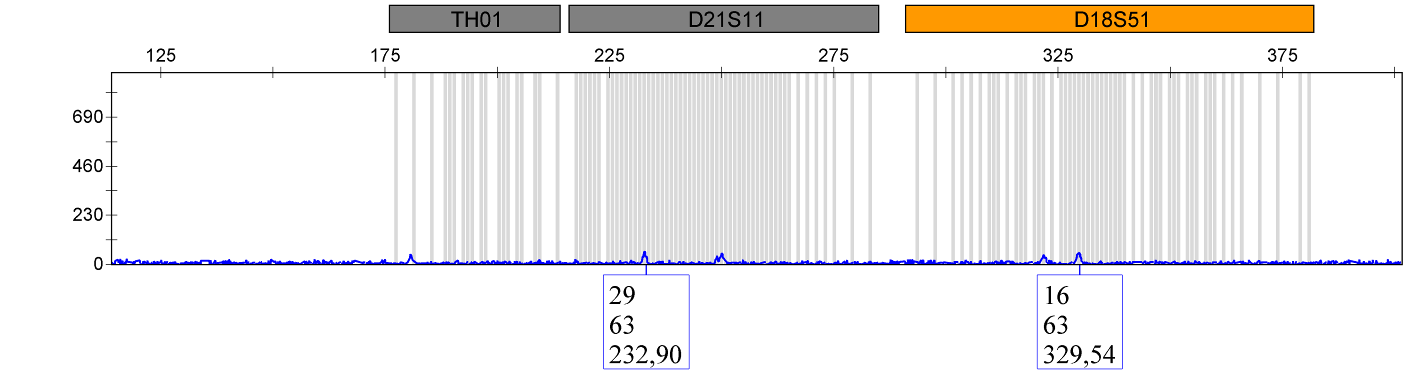

| | [redacted] RPA_panel | ✓ | ● |
|---|---|---|---|

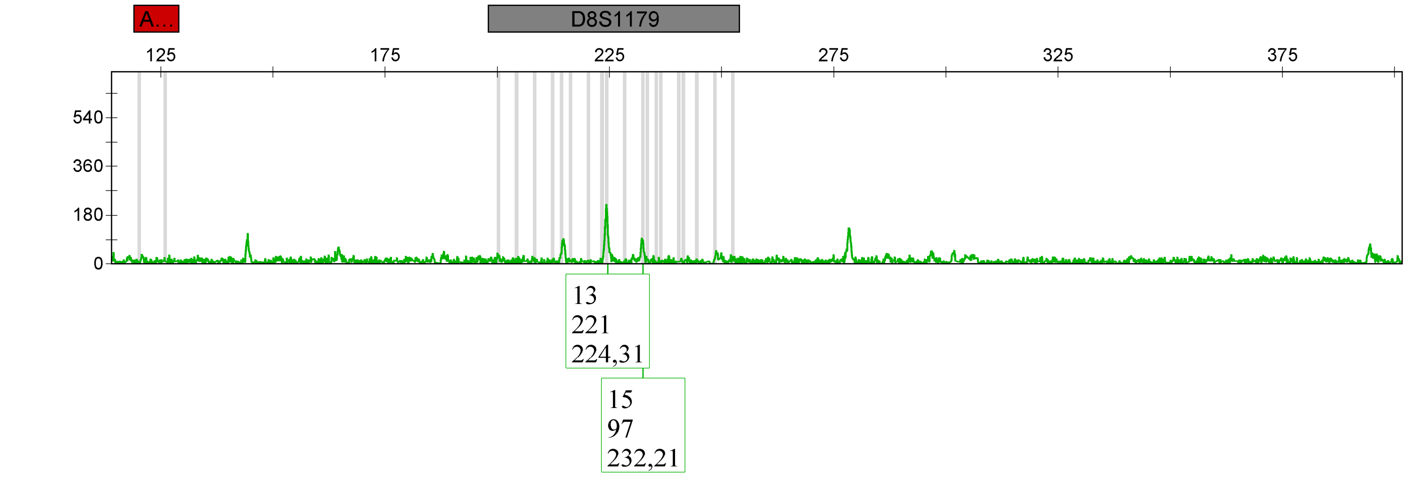

| Sample Name | Panel | SQO | CGQ |
|---|---|---|---|
| 500 ng replicate 2 | Sonja_RPA_panel | ✓ | |

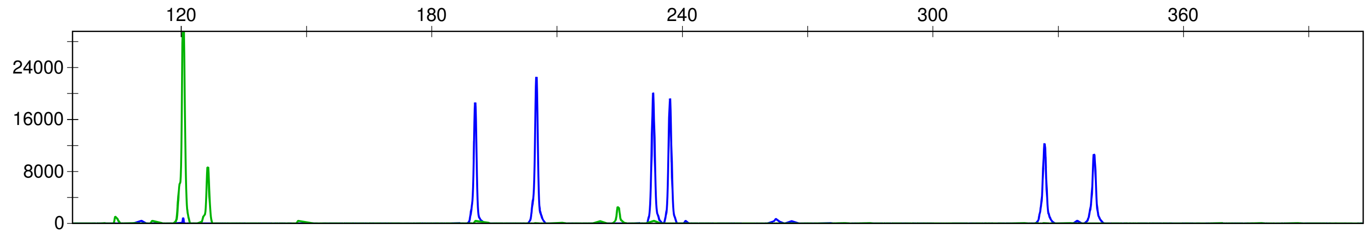

| Sample Name | Panel | SQO | CGQ |
|---|---|---|---|
| 250 ng replicate 2 | Sonja_RPA_panel | ✓ | |

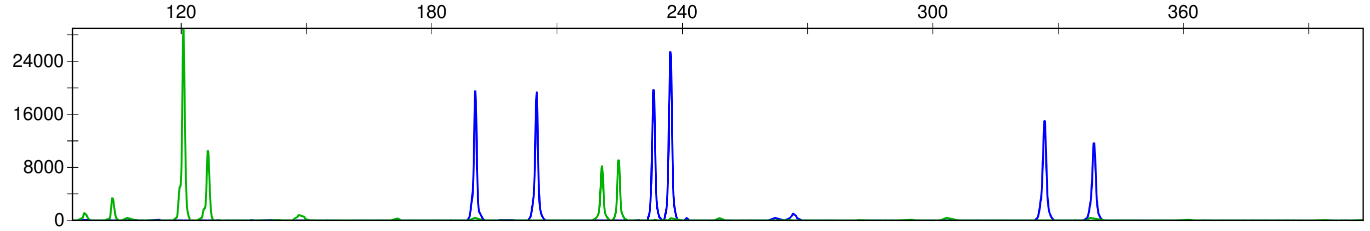

| Sample Name | Panel | SQO | CGQ |
|---|---|---|---|
| 125 ng replicate 2 | Sonja_RPA_panel | | |

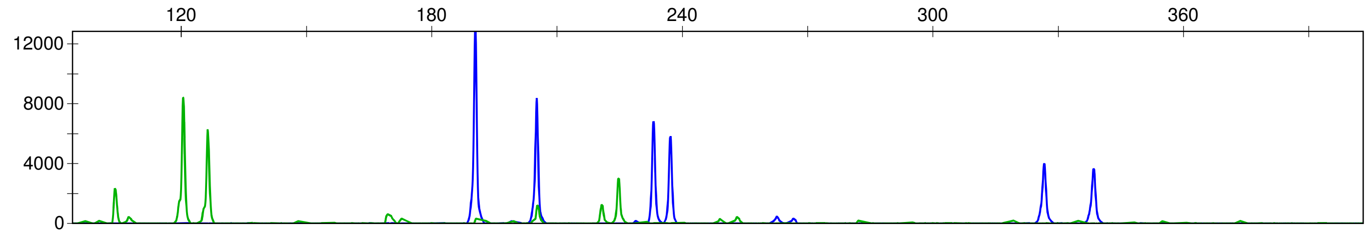

| Sample Name | Panel | SQO | CGQ |
|---|---|---|---|
| 62 ng replicate 2 | Sonja_RPA_panel | ✓ | |

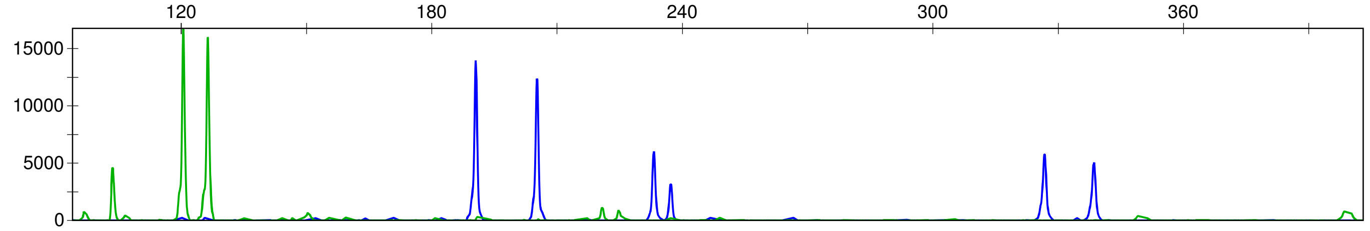

| Sample Name | Panel | SQO | CGQ |
|---|---|---|---|
| 31 ng replicate 2 | Sonja_RPA_panel | | |

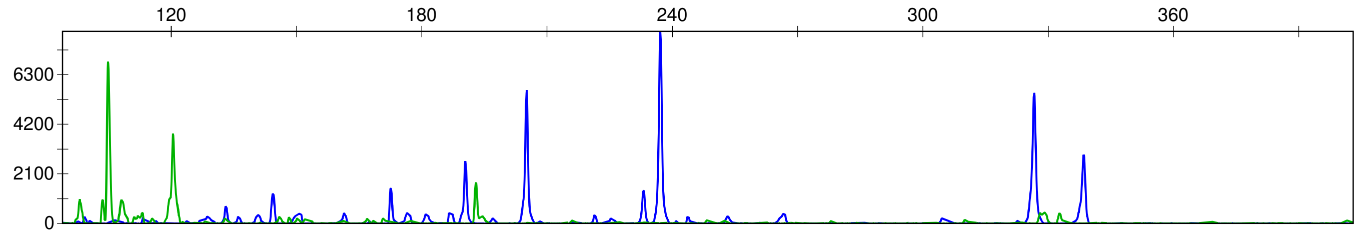
